# Exploring Inflammatory Dysregulation in Alveolar Macrophages: Implications for Novel Therapeutic Targets in Chronic Obstructive Pulmonary Disease

**DOI:** 10.1101/2024.03.20.585875

**Authors:** Saheed Adeyanju

## Abstract

Chronic obstructive pulmonary disease is a severe lung disease characterized by tissue destruction and limited airflow mainly caused by exposure to harmful environmental substances. Primary symptoms of this lung disorder include dyspnea, sputum production, and cough, which leads to respiratory failure. Prevalence increases with age, making it the most common cause of death worldwide. The primary objective of this study was to identify novel therapeutic targets via gene expression meta-analysis and to utilize them for drug reprofiling of FDA-approved drugs in treating chronic obstructive pulmonary disease. Multiple microarray and RNA-seq datasets from alveolar macrophages comprising healthy and diseased patients were processed to pinpoint significant dysregulated genes involved in this disease. Next, a meta-analysis was performed to identify the consistently differentially expressed genes in all datasets. Functional enrichment and protein-protein interaction analyses were conducted to single out the hub genes. Moreover, 3D structure prediction, virtual screening, and molecular dynamics simulations were utilized to explore the selected hub gene for drug repurposing. The number of significantly dysregulated genes identified via RNA-seq and microarray meta-analysis was found to be 104 and 57, respectively. Interestingly, VGLL3, ITIH5, ELOVL7, ACOD1, LAMB1, CXCL9, and GBP5 were common between the two sets revealing their significant association with the disease. CXCL9 and CCL3L3 were identified as the common hub genes between both sets. However, CXCL9, a chemokine, was prioritized for drug repurposing endeavors as it exhibits remarkable involvement in immune response and inflammation. Virtual screening of CXCL9 against selected drugs disclosed that CXCL9 has the highest binding affinity of −7.3 kcal/mol for Nintedanib, and binding affinities ranged from −2.4 kcal/mol to −7.3 kcal/mol. Moreover, Tepotinib and Crizotinib were found to be the second and third top-scoring drugs of −6.8 kcal/mol and −6.2 kcal/mol, respectively. Furthermore, the molecular dynamics simulation revealed that Crizotinib showed the most prominent results; however, its binding affinity is lower than Nintedanib. Therefore, Nintedanib is suggested as the better therapeutic agent to inhibit CXCL9 for treating chronic obstructive pulmonary disease.

**Highlights:**

- Meta-analysis of microarray and RNA-Seq datasets of alveolar macrophages from healthy and diseased patients disclosed novel therapeutic targets.
- Common significantly dysregulated hub gene CXCL9 is a novel drug target for COPD.
- CXCL9 is a chemokine responsible for inflammatory and immune responses utilized for drug reprofiling.
- Nintedanib, Tepotinib, and Crizotinib exhibited strong binding affinities against CXCL9.
- Virtual screening and simulation results revealed that inhibition of CXCL9 may be a potential treatment for COPD.

## 1. Introduction

Chronic obstructive pulmonary disease (COPD) is a chronic respiratory condition characterized by airflow limitation in the lungs [1]. It is a progressive disease that worsens over time. It is mainly caused by long-term exposure to irritating gases or particles, such as smoking cigarettes. Smoking is the primary risk factor for COPD, but air pollution, occupational dust and chemicals, and genetic factors can also contribute. COPD is a major cause of morbidity and mortality worldwide and is associated with significant healthcare costs [2].

COPD encompasses two main types: chronic bronchitis and emphysema. Former is characterized by inflammation and excessive mucus production in the bronchial tubes, causing a persistent cough and difficulty in clearing the airways [3]. On the other hand, emphysema involves the impairment of air sacs (alveoli) in the lungs, leading to reduced elasticity and decreased capacity for gas exchange. This results in shortness of breath and a sensation of breathlessness. COPD presents several symptoms, including a persistent cough (with or without sputum production), shortness of breath (especially during physical activity), wheezing, chest tightness, and frequent respiratory infections, among others [4].

There is currently no cure for COPD, and treatment focuses on symptom management and improving quality of life. Early diagnosis and proper management of COPD can help slow disease progression and alleviate symptoms [5]. Treatment commonly involves smoking cessation, medication usage, pulmonary rehabilitation, oxygen therapy, and, in advanced cases, surgical interventions [6]. However, COPD is a progressive disease, and despite treatment, it can continue to worsen over time. While treatment aims to slow down progression and manage symptoms, it cannot fully reverse lung damage. Not all individuals respond equally to available treatments, with some experiencing significant symptom improvement and others having limited response or requiring multiple treatment modalities [7]. Medications used for COPD management, such as bronchodilators and corticosteroids, can have side effects including increased heart rate, elevated blood pressure, oral thrush (with inhaled corticosteroids), and others [8]. It is important to balance the benefits of medication with potential side effects in COPD management.

One of the major issues with COPD is the economic burden on society. COPD imposes a significant economic burden on individuals, healthcare systems, and society. The costs associated with medications, hospitalizations, oxygen therapy, and other aspects of COPD management can be substantial [9].

Since, COPD is a multifactorial condition influenced by a combination of genetic predisposition, environmental exposures (such as smoking or air pollution), and individual susceptibility [10]. The pathogenesis of COPD involves intricate and incompletely understood mechanisms. Numerous molecular and genetic factors have been implicated in the development and progression of the disease. One notable example is Alpha-1 Antitrypsin Deficiency (AATD), a genetic disorder resulting from mutations in the SERPINA1 gene. These mutations lead to a deficiency in the alpha-1 antitrypsin (AAT) protein, which normally protects lung tissues from damage caused by enzymes like neutrophil elastase [11]. In the absence of sufficient AAT, uncontrolled activity of these enzymes occurs, accelerating the degradation of lung tissues and increasing the susceptibility to COPD, particularly among individuals who smoke [12].

Moreover, oxidative stress plays a significant role in COPD. Elevated oxidative stress occurs due to exposure to environmental factors such as cigarette smoke and air pollutants, which generate reactive oxygen species (ROS) capable of harming lung tissues [13]. An imbalance between oxidants and antioxidants in the lungs contributes to chronic inflammation, tissue injury, and the progression of COPD [14].

In addition, chronic inflammation serves as a hallmark of COPD, and various inflammatory mediators play a crucial role [15]. Cytokines, chemokines, and growth factors, among other molecules, contribute to the inflammatory response within the lungs. These molecules attract immune cells to the site of inflammation, leading to tissue damage and remodeling [16,17].

Alveolar macrophages, specialized immune cells residing in the lung alveoli, are pivotal in maintaining lung equilibrium, combating pathogens, and clearing respiratory system debris and foreign particles [18]. However, in the context of COPD, alterations in alveolar macrophage functionality can contribute to the development and progression of the disease, primarily through mechanisms involving inflammation, oxidative stress, and tissue damage [19].

The primary role of alveolar macrophages is to recognize and engulf pathogens, toxins, and other extraneous particles within the lungs. They release various inflammatory mediators, including cytokines and chemokines, to recruit other immune cells to the site of infection or injury [20]. In COPD, sustained exposure to cigarette smoke or other irritants can lead to persistent activation of alveolar macrophages, resulting in an exaggerated inflammatory response. This chronic inflammation can inflict damage on lung tissues, thereby fostering the development and progression of COPD [21].

Furthermore, alveolar macrophages produce ROS as part of their defense mechanism against pathogens. However, excessive ROS production or compromised antioxidant defenses can lead to oxidative stress, culminating in lung tissue damage [22]. In COPD, alveolar macrophages can generate an excessive amount of ROS due to prolonged exposure to cigarette smoke or environmental pollutants. This oxidative stress instigates inflammation, tissue injury, and the destruction of lung parenchyma, particularly observed in COPD-related emphysema [23].

The dysfunction of lung air sacs [alveoli) is a central feature in the impaired lung function observed in COPD. Therefore, investigating alveolar macrophages offers valuable insights into the underlying mechanisms of COPD pathogenesis and provides opportunities for the discovery of novel therapeutic biomarkers. Alveolar macrophages play a critical role in the release of diverse inflammatory mediators, such as cytokines, chemokines, and growth factors. These mediators actively contribute to the inflammatory response and subsequent tissue damage seen in COPD [24]. Profiling the gene expression of these inflammatory mediators allows for the identification of dysregulated molecular pathways that hold potential as biomarkers for disease diagnosis, monitoring disease progression, or evaluating treatment response [25,26].

This study aims to investigate gene dysregulation in COPD by analyzing multiple microarray and RNA-seq datasets from alveolar macrophages of COPD patients and healthy individuals. The integration of these datasets enabled a meta-analysis of differential expression, facilitating the identification of genes consistently dysregulated across all datasets. Functional enrichment analysis and protein-protein interaction analysis were then conducted on the dysregulated genes to identify hub genes. The hub genes identified in both microarray and RNA-seq studies were further explored as potential targets for drug repurposing. This involved utilizing 3D structure prediction, virtual screening, and molecular dynamics techniques to evaluate the compatibility of known drugs used for lung diseases with the hub gene. This knowledge can contribute to the development of more effective therapies and strategies for COPD patients.

## 2. Methodology

### 2.1 Overview of the Study

This study employed multiple microarray and RNA-seq datasets obtained from alveolar macrophages of individuals with COPD and compared them to those from healthy individuals. The primary objective was to identify genes that exhibit dysregulation in COPD. The integration of both microarray and RNA-seq data facilitated a meta-analysis differential expression analysis to identify genes that were dysregulated across all datasets. Subsequently, the dysregulated genes identified through meta-analysis were subjected to functional enrichment analysis, followed by protein-protein interaction analysis to identify hub genes. The hub genes that were commonly identified in both microarray and RNA-seq studies were then considered as potential targets for drug repurposing. This was achieved by employing 3D structure prediction, virtual screening, and molecular dynamics techniques to assess the inhibition compatibility of the known drugs used for lung diseases against the hub gene. The illustration of the study design is given in **Figure 1**.

**Figure 1:**
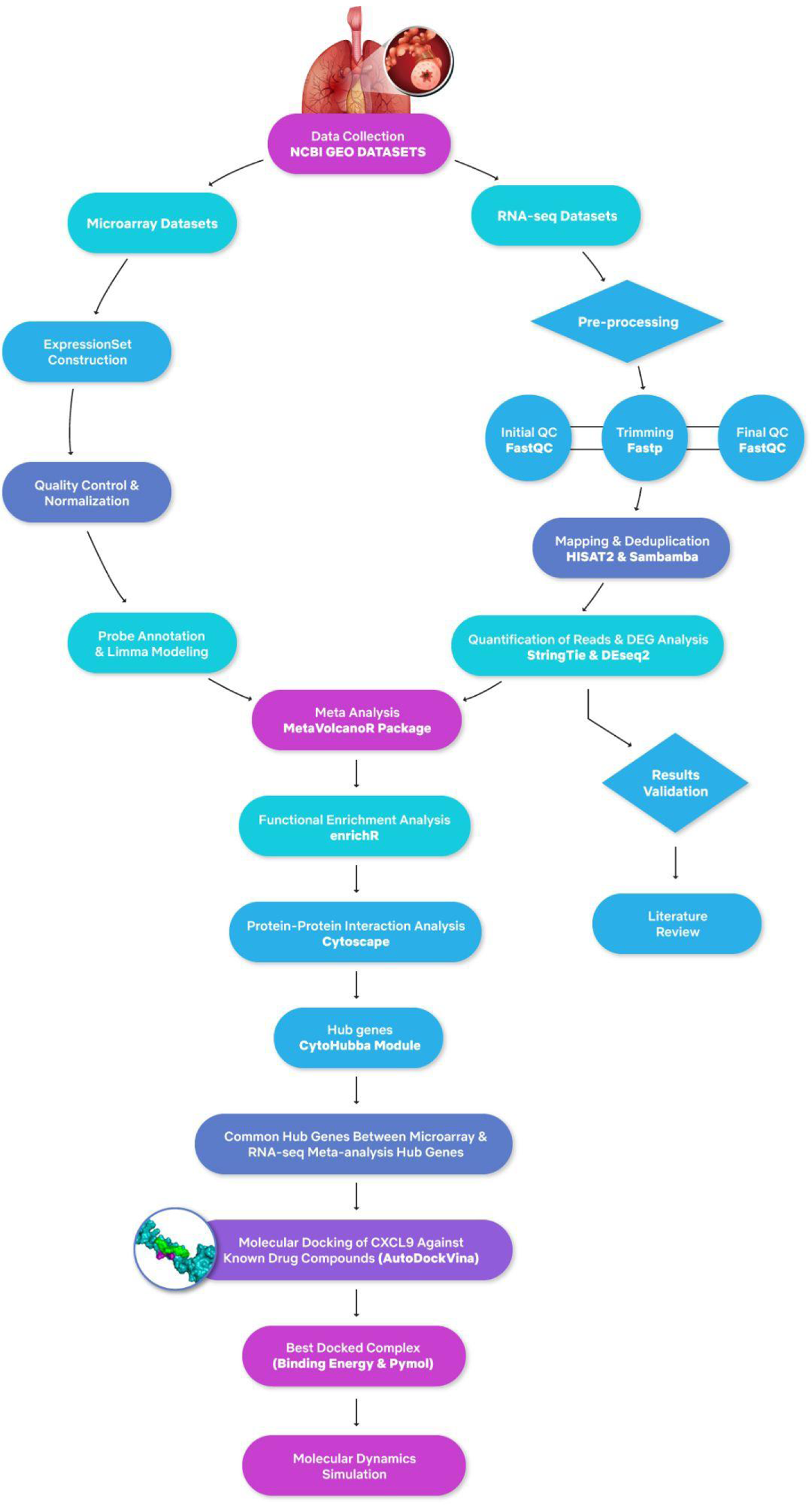
Overall workflow of COPD study

### 2.2 Data collection

The selection of datasets for this study was based on their relevance to COPD and specifically focused on alveolar macrophages obtained from individuals diagnosed with COPD, as well as samples from healthy individuals serving as a control group. A combined total of two RNA-seq datasets and four microarray datasets were gathered from the NCBI Gene Expression Omnibus (GEO) repository [27]. GEO serves as a global public repository encompassing functional genomics datasets, encompassing a wide range of high-throughput data such as micro-RNA profiles and next-generation sequencing data. The RNA-seq datasets utilized in this study include GSE183973 and GSE124180, while the microarray datasets employed are GSE16972, GSE130928, GSE112260, and GSE13896. The RNA-seq dataset with GEO ID GSE183973 was generated using the Illumina HiSeq 2000 platform. It comprises a total of 14 samples from individuals diagnosed with COPD and 14 samples from non-smoker healthy individuals. Furthermore, the RNA-seq dataset with GEO ID GSE124180 consists of 21 COPD cases and 4 samples from non-smoker individuals. The microarray dataset with GEO ID GSE112260 consists of 4 samples from individuals with COPD and 4 samples from non-smoker individuals. Additionally, the microarray dataset with GEO ID GSE130928 comprises 22 COPD samples and 24 samples from non-smoker healthy individuals.

### 2.3 RNA-seq preprocessing, mapping, and post-processing

To enhance the accuracy of subsequent analyses, preprocessing of the raw reads obtained from the sequencer was performed due to the presence of poor-quality reads, adapters, and primer contents. The FastQC tool was employed to assess multiple qualities of the raw reads, including per base sequence quality, GC content, per base N content, sequence length distribution, and levels of sequence duplication [28]. This preprocessing step aimed to improve the overall quality of the reads before further analysis. To eliminate poor-quality reads and contaminants, the FastP tool was utilized with specific parameters. The raw FASTQ sample was specified using the -i parameter, while the -o parameter was used to indicate the output file. Additionally, the -w parameter allowed for the utilization of multiple cores, enhancing processing efficiency. By employing FastP, PCR artifacts and low-quality reads were eliminated, thereby improving the overall data quality for subsequent analysis. Following the cleaning step, the FastQC tool was employed once more to evaluate the quality of the cleaned reads. Subsequently, the cleaned reads were aligned to the reference human genome (GRCh38) using the HISAT2 aligner [29]. To ensure accurate assessments of gene expression, duplicate reads were eliminated from the aligned reads. This was accomplished using the markdup module of Sambamba, a command-line tool, with the -r option enabled for appropriate adaptation [30]. By removing duplicates, the reliability of subsequent gene expression analyses was enhanced.

### 2.4 Read quantification and DEA analysis

The StringTie tool was used for measuring the level of gene expression [31]. The StringTie tool was employed to generate count reads through a series of three steps. Initially, it gathered alignments and constructed partial and full-length transcripts from millions of short-read sequences. Next, these transcripts were combined to create a consistent set of transcripts that could be compared across all samples. Finally, gene quantification was performed using the merged transcripts as a reference, and the -eB option was utilized to generate expression counts in Ballgown table format. For differential expression analysis between COPD patients and healthy controls, the DESeq2 tool was employed [32]. Biologically and statistically significant genes were identified based on FPKM-normalized expression values using log-fold change (logFC) values and p-values. Genes with a p-value less than 0.05 and a logFC value greater than 1 or less than −1 were considered as upregulated and dysregulated genes, respectively. To visualize the significant differentially expressed genes (DEGs), a Volcano plot was created. Additionally, the identification of upregulated and downregulated genes was validated by referring to relevant literature.

### 2.5 Microarray dataset analysis

The analysis of gene expression data was conducted using the R programming language. Subsequently, the Sample and Data Relationship Format (SDRF) file, which contains essential sample information, was read. The row names of the SDRF were assigned to the array data file column, resulting in the creation of an annotated data frame for further analysis. Specific columns representing the phenotype or condition of each sample were retained from the pData data frame. To facilitate subsequent analyses, a logarithmic transformation of the log2 expression data was performed, followed by the calculation of the ratio of the first two Principal Components (PCs) using Principal Component Analysis (PCA). Subsequently, the data underwent summarization, and background correction was performed without normalization. The rowMedians() function was employed to calculate the row medians of the expression values, while the sweep() function subtracted these row medians from each row of the expression values. Low-intensity transcripts were filtered out based on their intensity levels. Specifically, genes with median intensities below the predefined threshold of 4 were excluded from the analysis, while genes with median intensities above the threshold in at least the minimum required number of samples were retained. For gene annotation, the hgu133plus2.db package was utilized, enabling the annotation of genes with their corresponding symbols and names. In cases where probe IDs matched multiple genes, they were eliminated from the analysis. The limma package was then applied to all genes, with the creation of a contrast matrix to compare COPD and non-smoker individuals. The eBayes function was subsequently employed to calculate empirical Bayes moderated t-statistics, allowing for the identification of differentially expressed genes (DEGs). Notably, the subset function was employed to extract genes with a p-value less than 0.5, as well as logFC values less than −1 or greater than 1, thereby focusing on the most statistically significant and biologically relevant DEGs.

### 2.6 Meta-Analysis of RNA-seq and Microarray Gene Expression

To integrate and obtain a comprehensive understanding of gene expression values across multiple studies, separate meta-analyses were conducted for RNA-seq and microarray datasets. The R-based package MetaVolcanoR was utilized for performing the meta-analysis. The raw gene expression files derived from DESeq2 were used as input data for this analysis. The combining_mv function was called with various arguments to execute the meta-analysis. These arguments included the “diffexp” list, which encompassed the two datasets, criteria for selecting differentially expressed genes based on their p-values, column names for log2 fold change values and gene symbols, the method for combining fold changes across studies, and the significance threshold for the meta-analysis. Subsequently, the “MetaVolcano” function was employed to generate a volcano plot depicting the results of the meta-analysis. A significance threshold of p-value < 0.05 and logFC values <−1 and >1 were applied. By comparing the dysregulated genes identified from the meta-analyses of microarray and RNA-seq studies, common genes that exhibited either upregulation or downregulation in both types of studies were identified.

### 2.7 Functional Enrichment Analysis

To elucidate the molecular functions and biological pathways associated with the dysregulated and differentially expressed genes identified through meta-analysis, we employed the EnrichR 3.1 package. This particular package was specifically designed for the analysis of dysregulated genes and was executed within the widely used statistical programming language R 4.2.2 [33]. The upregulated and downregulated genes were analyzed separately to determine their respective functional enrichment. To visualize the results, bar plots of Gene Ontology (GO) terms and KEGG pathways were generated using the plotEnrich function of the genes. The enrichment terms were arranged based on their minimum p-values, allowing for a comprehensive representation of the most significant functional annotations associated with the dysregulated genes.

### 2.8 Protein-Protein Interactions and Identification of Hub Genes

The DEGs identified from both meta-analysis results were utilized as input in the STRING database, which encompasses protein-protein interaction (PPI) information derived from diverse sources, including databases, literature, and experimental validation. The PPIs were extracted from STRING using default parameters and subsequently employed as input in Cytoscape, a popular software platform for visualizing and analyzing biological networks. In Cytoscape, the CytoHubba module was utilized to identify the top 10 hub genes from the PPI network. Hub genes were determined based on their degree of centrality, which reflects the number of connections (edges) each gene has with other genes within the network. Genes with a higher degree of centrality are considered more central to the network, suggesting their potential importance in regulating biological processes related to the analyzed condition or disease. To identify common hub genes, the hub genes obtained from the meta-analysis-based DEGs of both microarray studies and RNA-seq studies were combined. By comparing the hub genes from each analysis, common genes that appeared as hub genes in both datasets were identified. These common hub genes were considered to have a higher level of significance and reliability, as their centrality in the PPI networks was consistently observed across different experimental techniques. The common hub gene (CXCL9) was then selected for further analysis, which is a chemokine that plays a role in inflammation, taking into consideration its potential functional importance in the context of the studied condition, COPD.

### 2.9 3D Structure Prediction and Selection of Drugs

In the absence of the complete 3D structure of CXCL9 (C-X-C motif chemokine ligand 9) on the Protein Data Bank, an alternative 3D structure available on AlphaFold (Q07325) was chosen for further analysis. The selection was based on the high confidence score of the AlphaFold prediction, specifically a pLDDT (predicted local distance difference test) score greater than 90, which indicates a high level of confidence in the predicted protein structure. For the CXCL9 protein, the compounds selected for further analysis were limited to approved drugs for various lung diseases from the Food and Drug Administration (FDA). By focusing on FDA-approved drugs, the study aimed to identify potential candidates for drug repurposing or exploration of their effects on CXCL9. These FDA-approved drugs have already undergone rigorous testing for safety and efficacy in various clinical indications, which makes them attractive candidates for further investigation. The rationale behind this approach is to leverage existing knowledge and resources to potentially find new therapeutic applications for FDA-approved drugs targeting CXCL9 for COPD treatment. The compounds were obtained from the DrugBank database [34] specifically selecting those with known potential for combating lung cancer [35] The rationale behind selecting these compounds is based on the assumption that they may also exhibit some effect on the CXCL9 protein. This assumption is supported by previous research indicating that CXCL9 protein is associated with an increased risk of lung cancer. By considering compounds that have demonstrated potential in treating lung cancer, there is a possibility that they could also modulate the activity or function of CXCL9, thus offering a potential avenue for therapeutic intervention in CXCL9-associated conditions, including COPD.

### 2.10 Molecular Docking and Visualization

The molecular docking of the CXCL9 protein was performed with the selected compounds that were retrieved from DrugBank [34]. Autodock Vina [36] was used for docking purposes because of its accuracy, flexibility, and efficiency. The protein (CXCL9) and the compounds were first prepared and saved in pdbqt format and config files were also created using the grid box. Subsequently, the docked complexes showed various binding affinities and among those complexes the best complex was selected which was then visualized on PyMOL (The PyMOL Molecular Graphics System, Version 2.0, Schrödinger, LLC). Different colors for protein and ligand were used in order to differentiate between them and the polar interactions were highlighted with their bond distances.

### 2.11 Molecular Dynamics Simulations

The molecular dynamics simulations of docked complexes of CXCL9 protein with the selected three compounds (Nintedanib, Tepotinib, and Crizotinib) were performed to analyze the comparative differences that resulted in the CXCL9 protein with each of the compounds. The simulations were executed using Maestro 12.0 (version 12.0.012, Schrödinger, LLC, New York, NY). The complexes were first prepared and optimized using the protein preparation wizard. Furthermore, a system builder was used to select the TIP3P water model and the structures were further minimized. Finally, the molecular dynamics simulations were performed at a temperature of 300K and a time period of 50 nanoseconds.

## 3. Results

### 3.1 Differential Expression Analysis and Meta-Analysis

MetaVolcanoR, a computational tool, was employed to conduct a meta-analysis of RNA-seq and four microarray studies individually. The objective was to identify genes that exhibited differential expression across both types of experiments. The meta-analysis of RNA-seq studies identified a total of 104 genes that displayed significant differential expression (Figure 2a). Among these genes, 78 were upregulated, indicating an increased expression level, while 26 were downregulated, indicating a decreased expression level. The top 20 upregulated and downregulated genes are given in **Table 1** and **Table 2**. In contrast, the meta-analysis of the microarray studies revealed 57 genes that exhibited significant differential expression. Among these genes, 18 were found to be upregulated, indicating an increased expression level, while 39 were downregulated, indicating a decreased expression level (Figure 2b). The top 20 upregulated and downregulated genes obtained from meta-analysis of microarray studies are given in **Table 3** and **Table 4**.

**Figure 2.**
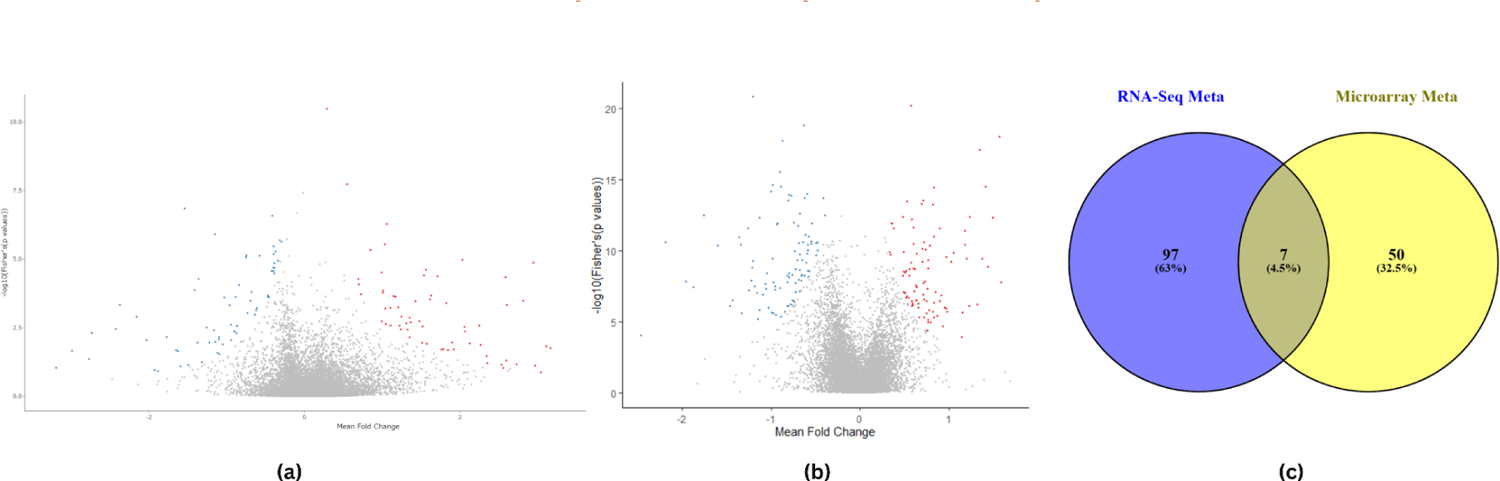
**(a)** Volcano plot for visualization of differentially expressed genes in RNA-seq meta-analysis studies, red color represents upregulated genes whereas blue color represents downregulated genes. **(b)** Volcano plot for visualization of differentially expressed genes in microarray meta-analysis studies, red color represents upregulated genes whereas blue color represents downregulated genes. **(c)** Venn diagram depicting the common genes between both meta-analysis studies of microarray and RNA-seq studies.

**Table 1.** Upregulated genes identified through meta-analysis in RNA-seq studies.

| SYMBOL | metap | metafc |
| --- | --- | --- |
| SIGLEC16 | 3.86E-09 | 4.828844377 |
| ENPP3 | 3.60E-05 | 3.808519658 |
| BPIFA1 | 0.01746667448 | 3.164076186 |
| ADAMTS1 | 0.0150137722 | 3.10864266 |
| STAP1 | 1.34E-05 | 2.943085627 |
| VGLL3 | 0.0003258687819 | 2.814389122 |
| ITIH5 | 0.0004641783831 | 2.602658703 |
| SCGB1A1 | 0.0488571429 | 2.597127367 |
| RNU1-28P | 4.53E-05 | 2.587268368 |
| LINC02104 | 0.03407901059 | 2.342321761 |
| NALCN-AS1 | 0.01334414115 | 2.266546924 |
| PDGFD | 0.002630386326 | 2.248881767 |
| RNA5SP217 | 0.01179263684 | 2.12751414 |
| GOLGA8A | 0.004308769031 | 2.064846555 |
| ELOVL7 | 0.002950183297 | 2.059030875 |
| HLA-DQA2 | 1.05E-05 | 2.031573066 |
| ACOD1 | 0.04741844162 | 1.974184701 |
| GBP7 | 0.01929058189 | 1.921154906 |
| RN7SL689P | 0.01092709072 | 1.900817113 |
| PTGER3 | 0.02064664513 | 1.847097809 |

**Table 2.** Downregulated genes identified through meta-analysis in RNA-seq studies.

| SYMBOL | metap | metafc |
| --- | --- | --- |
| PSPHP1 | 0.02167945214 | -2.986306909 |
| SOCS3 | 0.04361459238 | -2.766901128 |
| SLAMF9 | 0.004897533384 | -2.732654518 |
| CD1A | 0.003537009786 | -2.423943889 |
| C20orf96 | 0.0004651128633 | -2.370909116 |
| HLA-H | 0.001255641382 | -2.157439564 |
| ADGRG3 | 0.009035360782 | -2.026373084 |
| SEMA6B | 0.006850099929 | -1.770091695 |
| MMP1 | 0.02095514293 | -1.649298987 |
| UCHL1 | 0.02239481693 | -1.624745665 |
| TMEM216 | 1.42E-07 | -1.538729765 |
| FCER1A | 0.0001341971903 | -1.408465318 |
| RPL24P2 | 0.01107036761 | -1.374353339 |
| SLC26A11 | 5.21E-05 | -1.360421666 |
| LINC02470 | 0.003092902945 | -1.257643569 |
| FHL3 | 0.01104454219 | -1.220413992 |
| MVD | 0.003871158231 | -1.161908644 |
| HLA-DRB5 | 1.22E-06 | -1.150595474 |
| CD1C | 0.01045544644 | -1.145244942 |
| ZNF888-AS1 | 0.02570760693 | -1.117270857 |

**Table 3.** Upregulated genes identified through meta-analysis in microarray studies.

| **SYMBOL** | **metap** | **metafc** |
| --- | --- | --- |
| SNAR-D | 0.03712458443 | 1.633370889 |
| AFAP1L1 | 1.72E-08 | 1.58988499 |
| FBXO15 | 1.04E-18 | 1.569427323 |
| SLC39A10 | 5.50E-13 | 1.494567543 |
| TMEM163 | 1.51E-09 | 1.44295042 |
| SRPX | 3.65E-15 | 1.411955638 |
| TRERF1 | 3.84E-10 | 1.373053738 |
| SH3RF1 | 9.28E-18 | 1.352961782 |
| GPR34 | 6.45E-07 | 1.319083943 |
| SMIM3 | 4.76E-13 | 1.237436264 |
| TDRD9 | 8.12E-07 | 1.227086543 |
| CNIH3 | 3.67E-10 | 1.200532375 |
| DNASE2B | 4.53E-12 | 1.181098139 |
| DDIT4L | 4.27E-11 | 1.180550631 |
| GPNMB | 2.31E-06 | 1.155869397 |
| SLC18B1 | 0.0001254955234 | 1.145926946 |
| MLLT11 | 1.52E-10 | 1.053148525 |
| SPRY2 | 6.22E-10 | 1.024412139 |
| VGLL3 | 7.62E-15 | -1.000673278 |
| PSTPIP2 | 1.04E-06 | -1.024928124 |

**Table 4.** Downregulated genes identified through meta-analysis in microarray studies.

| **SYMBOL** | **metap** | **metafc** |
| --- | --- | --- |
| LINC02244 | 9.56E-05 | -2.466298748 |
| RSPO3 | 2.88E-11 | -2.182790429 |
| SHROOM3 | 1.49E-08 | -1.958524487 |
| ELOVL7 | 4.17E-08 | -1.868868228 |
| SERPINB2 | 0.004765856933 | -1.747789267 |
| GCH1 | 4.44E-11 | -1.593699629 |
| TNFAIP6 | 8.62E-07 | -1.458178849 |
| CCL23 | 3.33E-07 | -1.428746711 |
| LAMB1 | 1.32E-11 | -1.35662363 |
| TFPI2 | 0.00293524751 | -1.351216903 |
| IL32 | 3.99E-11 | -1.336058056 |
| ANKRD22 | 2.61E-06 | -1.334363853 |
| PDE4B | 2.83E-12 | -1.260759394 |
| ACOD1 | 6.45E-08 | -1.243767762 |
| SCARNA5 | 0.04552494833 | -1.234004436 |
| RCAN2 | 1.31E-08 | -1.220584742 |
| NDP | 0.0002741341478 | -1.214632617 |
| MCOLN2 | 1.70E-09 | -1.209832014 |
| LPAR3 | 1.64E-21 | -1.204327707 |
| HLA-DQB2 | 6.13E-10 | -1.199847533 |

Remarkably, a subset of 7 (VGLL3, ITIH5, ELOVL7, ACOD1, LAMB1, CXCL9, GBP5) genes was identified as being commonly differentially expressed in both meta-analysis of RNA-seq and microarray studies (Figure 2c). This finding highlights the presence of shared genes that are consistently dysregulated across different experimental platforms, providing valuable insights into the molecular alterations associated with COPD. The overlap between the two types of studies strengthens the robustness and reliability of the identified differentially expressed genes. Further analysis of these commonly expressed genes may uncover key biological pathways and molecular mechanisms underlying COPD. The identified DEGs from each study individually are given in **Supplementary Sheet 1**.

### 3.2 Functional Enrichment Analysis of Meta-Analysis-based Dysregulated Genes

To gain a deeper understanding of the functional implications of the dysregulated genes identified through meta-analysis, a functional enrichment analysis was performed. This analysis helps identify the biological processes, molecular functions, and cellular components that are significantly enriched among the dysregulated genes. The functional enrichment analysis provides insights into the potential roles and interactions of these genes within biological pathways and cellular processes.

The GO enrichment analysis of the dysregulated genes obtained from RNA-seq meta-analysis revealed several interesting findings. The upregulated genes were found to be associated with molecular functions such as CXCR3 and CXCR chemokine receptor binding, fatty-acyl-CoA synthase activity, C-acyltransferase activity, and CoA-ligase activity. This suggests their involvement in chemokine receptor signaling, lipid metabolism, and cellular energy processes **Supplementary Document 1 Figure S1**. In contrast, the downregulated genes showed a different pattern of molecular functions, including RNA and rRNA binding, oxidoreduction-driven active transmembrane transport, NADH dehydrogenase activity, ubiquitin ligase inhibitor activity, and ubiquitin-protein ligase inhibitor activity **Supplementary Document 1 Figure S2**. These findings indicate a potential disruption in RNA processing, cellular respiration, and protein degradation pathways. GO cellular component analysis revealed that the upregulated genes were predominantly expressed in intracellular vesicles, membrane attack complexes, and HFE-transferrin receptor complexes in **Supplementary Document 1 Figure S3**. This suggests their involvement in intracellular transport and immune-related processes. On the other hand, the downregulated genes were found to be expressed in cytosolic large and small ribosomal subunits, ribosomes, polysomal ribosomes, and mitochondrial membranes, indicating a potential impact on protein synthesis and mitochondrial function **Supplementary Document 1 Figure S4**. In terms of biological processes, the upregulated genes were significantly associated with adenylate cyclase-activating G protein signaling, cellular response to lipopolysaccharides, and cellular response to interferon-gamma **Supplementary Document 1 Figure S5**. These findings suggest their involvement in immune responses and signaling pathways related to inflammation and antiviral defense. In contrast, the downregulated genes were found to be involved in various processes, including protein targeting to the endoplasmic reticulum, cotranslational protein targeting to the membrane, cytoplasmic translation, nuclear-transcribed mRNA catabolic process, peptide biosynthetic process, and rRNA processing **Supplementary Document 1 Figure S6**. These findings imply potential disruptions in protein synthesis, protein trafficking, and mRNA metabolism. Furthermore, the pathway analysis revealed the upregulation of pathways such as Toll-like receptor signaling, butanoate metabolism, cytokine-cytokine receptor interaction, and chemokine signaling **Supplementary Document 1 Figure S7**. These pathways are closely related to immune responses, inflammation, and cellular signaling processes. In contrast, the downregulated pathways included ribosome and oxidative phosphorylation, indicating potential alterations in protein synthesis and cellular energy metabolism **Supplementary Document 1 Figure S8**.

The GO enrichment analysis of the dysregulated genes obtained from microarray meta-analysis revealed interesting findings. The upregulated genes were associated with molecular functions such as syndecan binding, endodeoxyribonuclease activity, G protein-coupled purinergic nucleotide binding, MAP-kinase scaffold activity, and purinergic nucleotide receptor activity **Supplementary Document 1 Figure S9**. These functions suggest their involvement in extracellular matrix interactions, nucleotide metabolism, and intracellular signaling processes. In contrast, the downregulated genes showed different molecular functions, including serine-type endopeptidase activity, cAMP-dependent protein kinase inhibitor and regulator activity, and phosphatidylinositol phosphate binding **Supplementary Document 1 Figure S10**. These findings indicate potential alterations in protease activity, protein kinase regulation, and lipid signaling pathways. GO cellular component analysis revealed that the upregulated genes were predominantly expressed in piP-body, integral components of synaptic vesicles, intrinsic components of synaptic vesicles, P granules, and AMPA glutamate receptor complexes **Supplementary Document 1 Figure S11**. These findings suggest their involvement in neuronal signaling and RNA processing. On the other hand, the downregulated genes were found to be expressed in integral components of the lumenal side of the endoplasmic reticulum, the luminal side of the endoplasmic reticulum, ER to Golgi transport vesicle membranes, coated vesicle membranes, and mitochondrial envelopes **Supplementary Document 1 Figure S12**. This indicates potential disruptions in protein synthesis, intracellular trafficking, and mitochondrial function. In terms of biological processes, the upregulated genes were significantly associated with positive regulation of mitochondrial depolarization and negative regulation of lens fiber cell differentiation **Supplementary Document 1 Figure S13**. These findings suggest their involvement in mitochondrial function and lens development. In contrast, the downregulated genes were found to be involved in various processes, including cytokine-mediated signaling pathways, cellular response to lipopolysaccharides, cellular response to bacteria, and positive regulation of cytokine production **Supplementary Document 1 Figure S14**. These findings indicate potential alterations in immune responses and inflammatory processes. Interestingly, only one pathway, namely lysosome, was found to be highly upregulated **Supplementary Document 1 Figure S15**. This suggests potential dysregulation of lysosomal function and degradation pathways. On the other hand, several pathways, including Th1 and Th2 cellular differentiation, Th17 cell differentiation, C-type lectin receptor signaling pathway, osteoclast differentiation, and MAPK signaling pathway, were found to be downregulated **Supplementary Document 1 Figure S16**. These pathways are involved in immune responses, cell differentiation, and intracellular signaling.

### 3.3 Protein-Protein Interactions and Hub Genes Identification

Through the STRING database, PPIs of the dysregulated genes obtained from the meta-analysis of both microarray and RNA-seq studies were identified separately. These PPIs provide valuable information about the functional relationships and potential interactions among the dysregulated genes. The network analyses of the dysregulated genes from RNA-seq meta-analysis and microarray meta-analysis, respectively are presented in **Supplementary Document 1** as **Figure S17** and **Figure S18**. These network visualizations illustrate the connections and interactions between the dysregulated genes, highlighting their complex relationships within COPD. Furthermore, based on the network analysis results, the top 10 hub genes were identified for each meta-analysis, as shown in **Table 5** and **Table 6** for RNA-seq and microarray meta-analysis, respectively, and in Figure 3a **and 3b**, respectively. These hub genes play crucial roles in the network and are likely to have significant functional and regulatory implications. Interestingly, the common genes CXCL9 and CCL3L3 were identified in both the RNA-seq and microarray meta-analysis. However, considering its elevated degree of centrality, significant involvement in immune response mechanisms, and extensive interactions with other genes within the network, CXCL9 was prioritized for drug repurposing endeavors. CXCL9, classified as a chemokine, assumes a critical function in orchestrating immune responses and inflammatory processes. Its primary role revolves around the recruitment and activation of immune cells, notably T cells, at sites of infection or inflammation.

**Figure 3:**
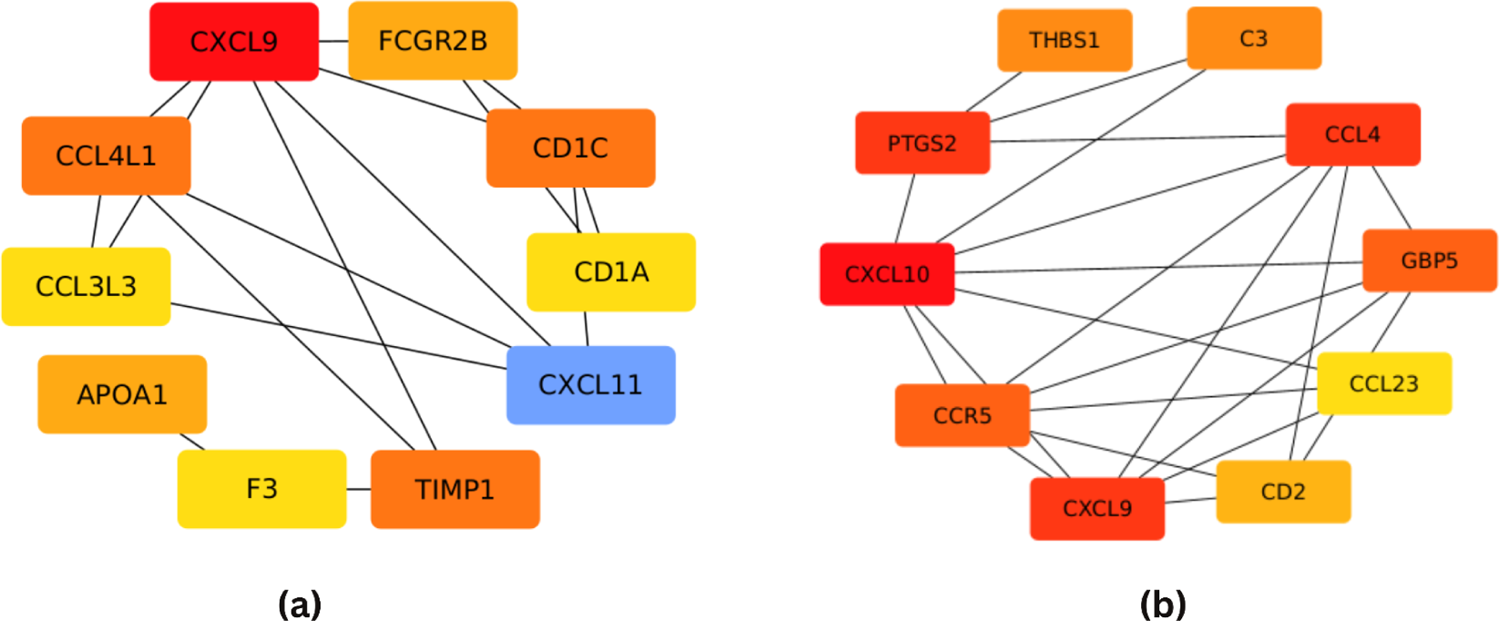
**(a)** Top 10 hub genes in RNA-seq-based meta-analysis dysregulated genes identified through protein-protein interaction network analysis. The darker color represents more interactions within the same network. **(b)** Top 10 hub genes in microarray-based meta-analysis dysregulated genes identified through protein-protein interaction network analysis. The darker color represents more interactions within the same network.

**Table 5.** Top 10 hub genes in RNA-seq-based meta-analysis genes and their score obtained through CytoHubba in Cytoscape.

| Name | Score |
| --- | --- |
| CXCL9 | 7 |
| CXCL11 | 6 |
| TIMP1 | 5 |
| CCL4L1 | 5 |
| CD1C | 5 |
| FCGR2B | 4 |
| APOA1 | 4 |
| CD1A | 3 |
| CCL3L3 | 3 |
| F3 | 3 |

**Table 6.** Top 10 hub genes in microarray-based meta-analysis genes and their score obtained through CytoHubba in Cytoscape.

| Name | Score |
| --- | --- |
| CXCL10 | 10 |
| CXCL9 | 7 |
| CCL4 | 7 |
| PTGS2 | 7 |
| CCR5 | 6 |
| GBP5 | 6 |
| C3 | 5 |
| THBS1 | 5 |
| CD2 | 4 |
| CCL23 | 3 |

### 3.4 3D Structure Prediction and Selection of Drugs

An alternative structure for CXCL9 protein which was obtained from the AlphaFold (Q07325), is shown in Figure 4a. Furthermore, the compounds (**Table 7**) were retrieved from the DrugBank [34] that already had the potential to fight against lung cancer which is the reason the selected compounds were thought to have some effect on the CXCL9 protein as well, as the CXCL9 protein is also associated with a risk of lung cancer [35].

**Figure 8.**
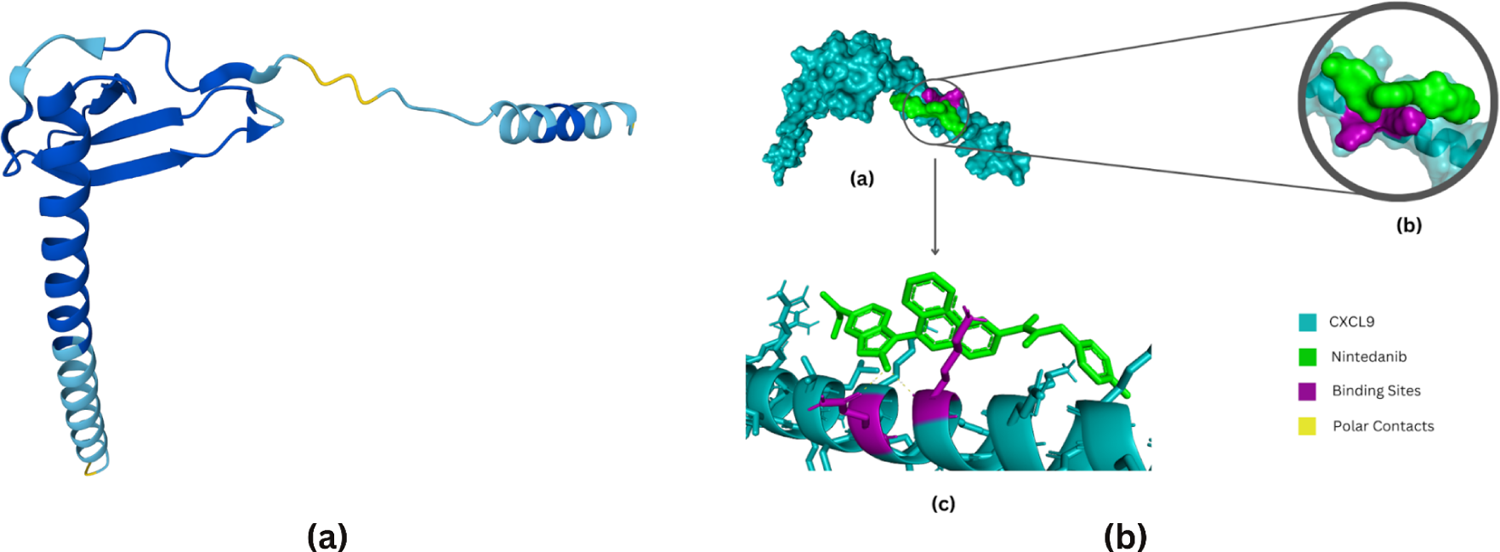
**(a)** C-X-C motif chemokine 9 (CXCL9) 3D protein structure obtained from AlphaFold. **(b)** Visualization of docked CXCL9 protein with Nintedanib on PyMOL.

**Table 7.**
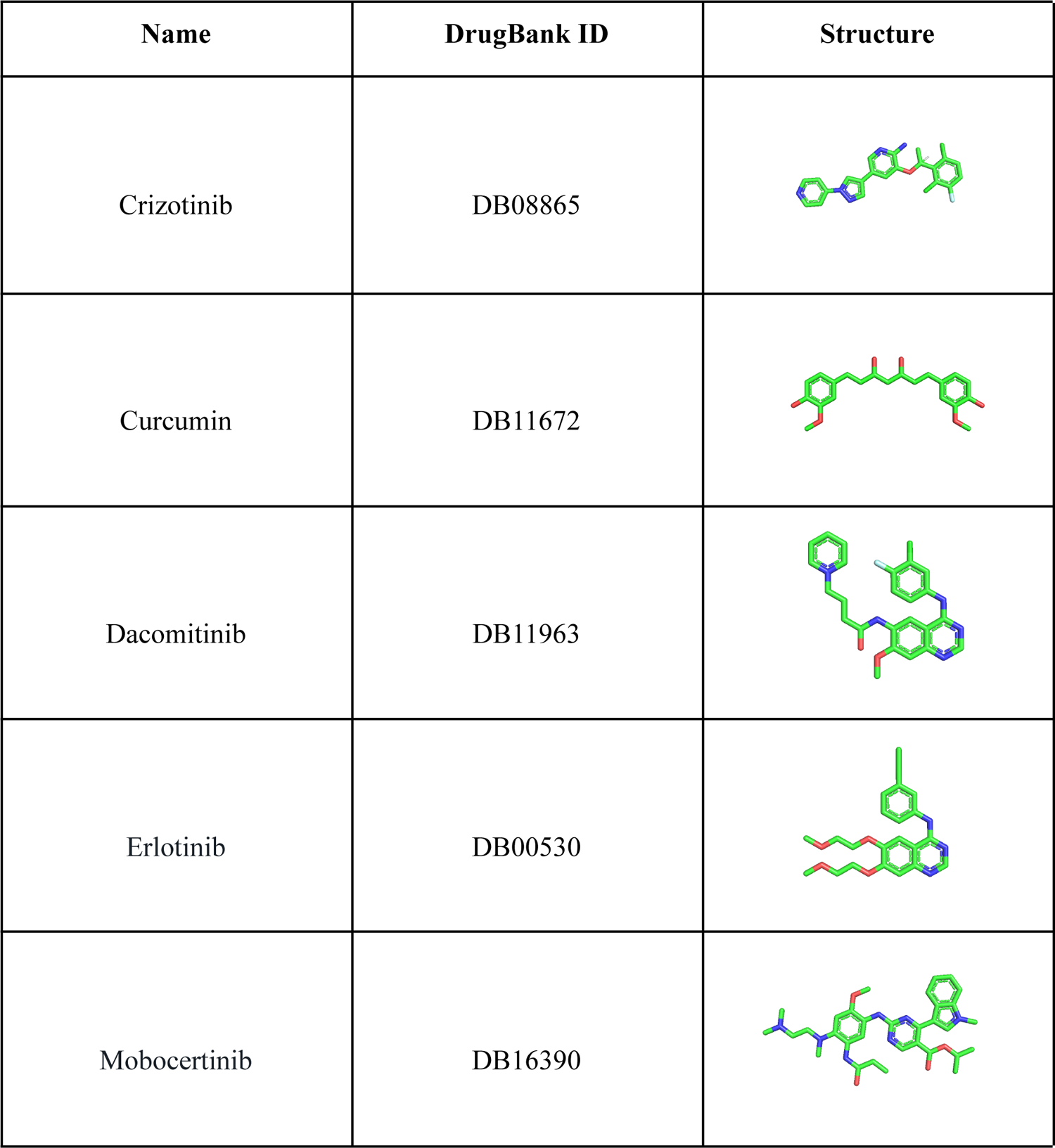
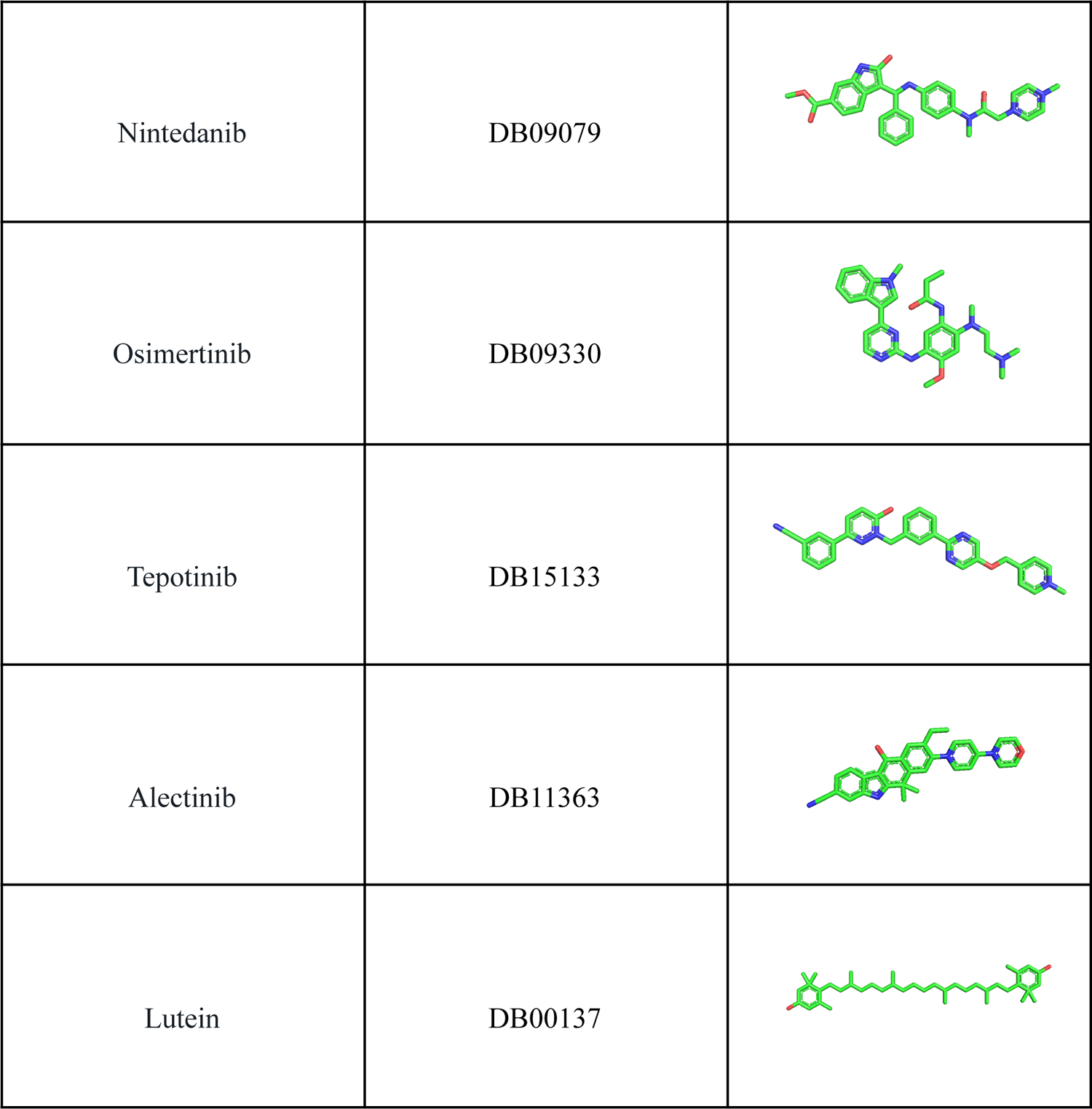
Drug compounds with their names, DrugBank IDs and structures.

### 3.5 Molecular Docking

The molecular docking was performed for CXCL9 with retrieved compounds using AutoDock Vina and resulted in binding affinities that ranged from −2.4 (kcal/mol) to −7.3 (kcal/mol) (**Table 8**). The first three best complexes based on the binding affinities were of CXCL9-Nintedanib, CXCL9-Tepotinib, and CXCL9-Crizotinib, having the binding affinities of −7.3 (kcal/mol), −6.8 (kcal/mol) and −6.2 (kcal/mol), respectively. The CXCL9-Nintedanib complex interactions are shown in Figure 4b.

**Table 8.** Docked complexes of compounds with CXCL9 and their binding affinities.

| Compounds Docked with CXCL9 | Binding Affinities (kcal/mol) |
| --- | --- |
| Nintedanib (DB09079) | -7.3 |
| Tepotinib (DB15133) | -6.8 |
| Crizotinib (DB08865) | -6.2 |
| Alcetinib (DB11363) | -5.6 |
| Erlotinib (DB00530) | -5.6 |
| Osimertinib (DB09330) | -5.6 |
| Dacomitinib (DB11963) | -4.9 |
| Mobocertinib (DB16390) | -4.8 |
| Curcumin (DB11672) | -4.5 |
| Lutein (DB00137) | -2.4 |

### 3.6 Molecular Dynamics Simulations

The molecular dynamics simulation results of CXCL9-Nintedanib complex indicated that the protein showed stability from 34.65 ns to the end of the simulation time, while it fluctuated from the beginning of the simulation to 32.35 ns time, 21.11 Å being the highest fluctuation recorded at 29.35 ns period. It was found to be overall unstable as it fluctuated considerably. On the other hand, the ligand fluctuated from 5.45 ns to 20.90 ns time and 34.30 ns to 47.55 ns time. The highest fluctuation was 21.74 Å at 43.55 ns time and it was also found to be overall unstable. The minimum difference between the protein and the ligand was at 2.95 ns period, while the RMSD values of the protein and the ligand at the end of the simulation time were 16.93 Å and 9.02 Å, respectively (Fig 5a). Furthermore, the protein-RMSF plot showed that the protein residues fluctuated significantly at positions 1-48,50-62,66-73, and 77-125. The highest fluctuations were shown by MET (16.27 Å) at position 1, followed by LYS (15.08 Å) at position 2, LYS (13.45 Å) at position 3, PRO (13.08 Å) at position 56, ASN (12.75 Å) at position 69, SER (12.38 Å) at position 55, GLY (11.87 Å) at position 70, and SER (11.46 Å) at position 4, while other protein residues at positions 5-29,31-48,50-54,57-62,66-68,71-73, and 77-125 fluctuated considerably more than 5 Å respectively (Fig 5b). Furthermore, the protein-ligand contacts indicated that LYS-100 exhibited the maximum interactions (Hydrogen Bonds = 0.036, Hydrophobic = 0.295, Water Bridges = 0.115), followed by LYS-103 (Hydrophobic = 0.247, Water Bridges = 0.166), LYS-104 (Hydrophobic = 0.006, Water Bridges = 0.335), GLN-99 (Hydrogen Bonds = 0.054, Water Bridges = 0.149), and GLY-102 (Hydrogen Bonds =0.097, Water Bridges = 0.040) (Fig 6a). At last, the ligand-protein contacts showed that LYS-104 interacted directly with the ligand molecule for 33% of the simulation time (Fig 6b).

**Figure 5.**
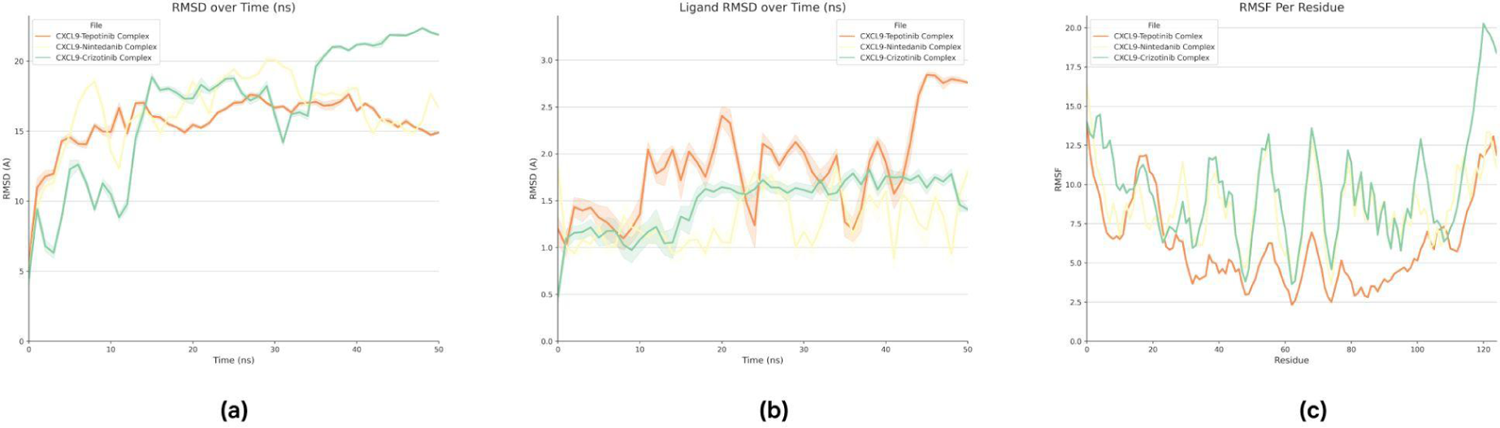
Molecular dynamics simulation results of CXCL9 protein complexes with Nintedanib, Tepotinib and Crizotinib performed on Maestro. Protein-ligand RMSD (a),(b). Protein-RMSF (c).

**Figure 6.**
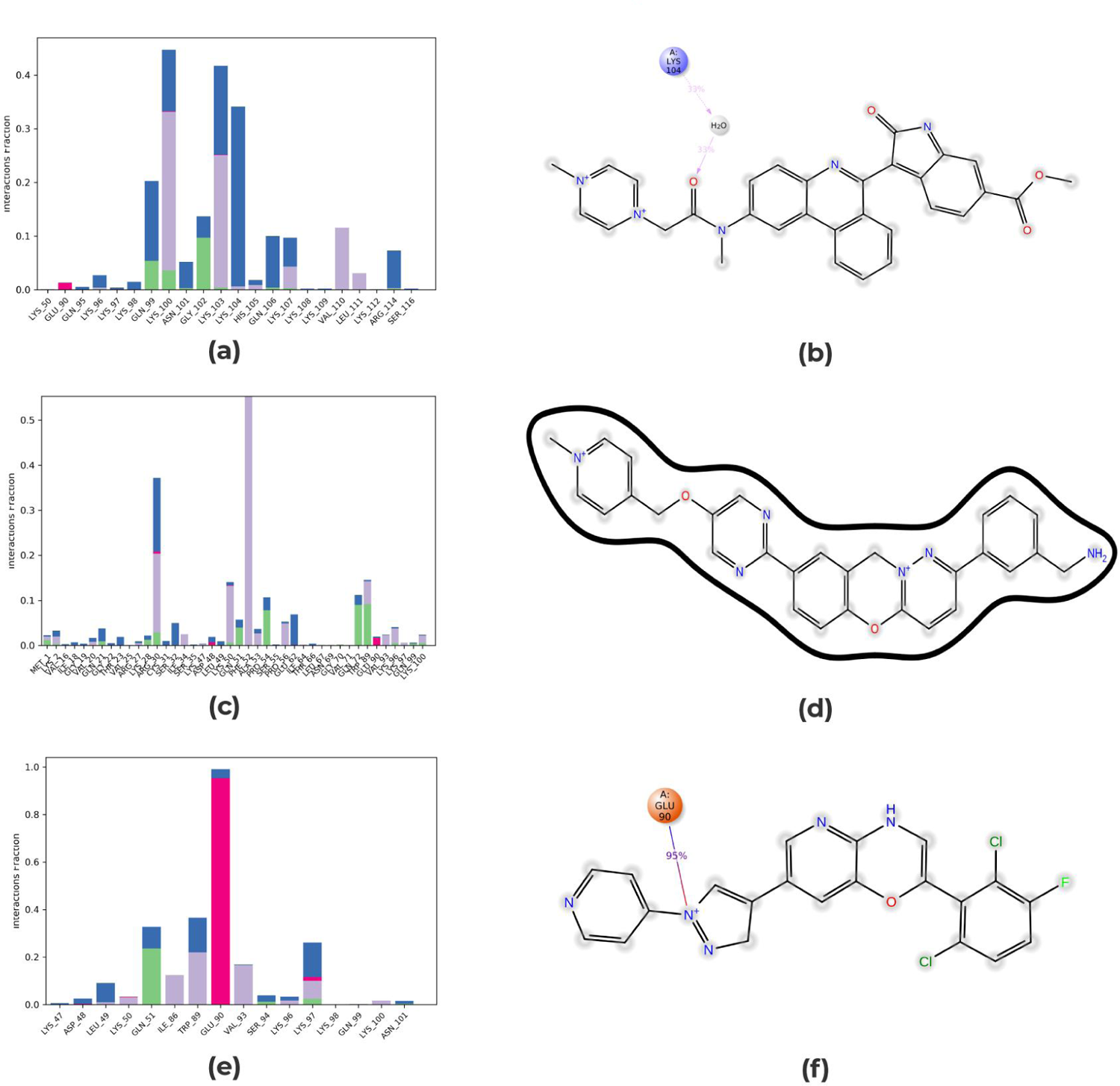
Molecular dynamics simulation results of CXCL9 protein complexes with Nintedanib, Tepotinib and Crizotinib performed on Maestro. Protein-ligand contacts (a), (c), (e). Ligand-protein contacts (b), (d), (f).

The simulation results of the CXCL9-Tepotinib complex indicated that the protein was stable from 15.00 ns to 22.95 ns time, while it fluctuated from the beginning of the simulation to 14.80 ns and 37.35 ns to 39.95 ns time. It showed the highest fluctuation of 18.42 Å at 39.55 ns period, and was found to be overall unstable. Instead, the ligand fluctuated from the beginning of the simulation to 11.60 ns time. It showed the highest fluctuation of 33.56 Å at 10.95 ns time and was found to be overall unstable. The minimum difference between the protein and the ligand was found at 0.20 ns time, while the RMSD values of the protein and the ligand at the end of the simulation time were 14.78 Å and 26.57 Å, respectively (Fig 5c). Moreover, the protein-RMSF plot indicated considerable protein residue fluctuations at positions 1-30,39,44,45,54-58,68-71,78, and 100-125. The highest fluctuation was of MET (13.95 Å) at position 1, afterwards THR (13.35 Å) at position 124, LYS (12.51 Å) at positions 2 and 122, THR (12.25 Å) at position 125, LYS (12.21 Å) at position 123, and GLN (11.87 Å) at position 121, while other protein residues at positions 3-30, 39,44,45,54-58,68-71,78, and 100-120 showed considerable fluctuations of more than 5 Å, respectively (Fig 5d). Furthermore, the protein-ligand contacts showed that PHE-52 exhibited the maximum interactions (Hydrophobic = 0.553), followed by ARG-30 (Hydrogen Bonds = 0.029, Hydrophobic = 0.175, Ionic = 0.005, Water Bridges = 0.164), TRP-89 (Hydrogen = 0.092, Hydrophobic = 0.051, Water Bridges = 0.003) (Fig 6c). Finally, the ligand-protein contacts showed that there were no protein residues that interacted with the ligand molecule for more than 30% of the simulation time (Fig 6d).

The simulation results of the CXCL9-Crizotinib complex indicated that the protein showed stability from 6.85 ns to 8.10 ns time, while it fluctuated significantly from 13.00 ns to 30.10 ns time and from 34.85 ns to the end of the simulation time. It showed the highest fluctuation of 22.79 Å at 48.80 ns time and was found to be overall unstable. On the other hand, the ligand fluctuated from the beginning of the simulation to 25.70 ns time and from 35.40 ns to 45.90 ns period. It showed stability from 30.45 ns to 34.75 ns time but was found to be overall unstable. The minimum difference between the protein and the ligand was recorded at 1.45 ns time, while the RMSD values of the protein and the ligand at the end of the simulation time were 21.96 Å and 10.05 Å, respectively (Fig 5e). Furthermore, the protein-RMSF plot showed that the protein residues significantly fluctuated at positions 1-47,50-61,66-74, and 76-125. The highest fluctuating residue was LYS (20.27 Å) at position 122, followed by GLN (20.23 Å) at position 121, THR (19.41 Å) at position 124, LYS (19.04 Å) at position 123, and THR (18.07 Å) at position 125, while other protein residues at positions 1-47,50-61,66-74, and 76-120 were also considerably fluctuating with RMSF value of more than 5 Å respectively (Fig 5f). Moreover, the protein-ligand contacts showed that GLU-90 exhibited the maximum interactions (Ionic = 0.952, Water Bridges = 0.038), followed by TRP-89 (Hydrophobic = 0.218, Water Bridges = 0.147), and GLN-51 (Hydrogen Bonds = 0.237, Water Bridges = 0.091) (Fig 6e). At last, the ligand-protein contacts indicated that GLU-90 interacted with the ligand molecule for 95% of the simulation time (Fig 6f).

## 4. Discussion

COPD, encompassing emphysema and chronic bronchitis, is the third leading cause of morbidity and mortality worldwide. The objective of this study was to identify dysregulated genes associated with COPD by conducting a meta-analysis of RNA-seq and microarray datasets. The datasets were obtained from COPD-positive and healthy individuals, specifically focusing on alveolar macrophages of the lung. The RNA-seq datasets used in this study were GSE124180 and GSE183973. DGE analysis of GSE124180 identified a total of 168 dysregulated genes, including 130 upregulated genes and 38 downregulated genes. Similarly, DGE analysis of GSE183973 predicted a total of 609 dysregulated genes, with 390 genes upregulated and 219 genes downregulated. Notably, 36 genes were found to be dysregulated in both datasets, namely CAMP, CCL3L3, CCL4L2, ITGBB, ABCA1, PDGFD, CXCL2, TNAFIP6, GOS2, PTGS2, CCL20, ACOD1, IGF1, FGF10, DDX43, CADPS2, FLT1, FOLR3, ENPP5, ITIH5, CXCL9, SHROOM3, CXADR, ADAMTS1, CD1A, CD1B, ELOVL7, CACNB3, CXCL11, ITSN1, VGLL3, RDUR, NALCN-AS1, TENM2-AS1, LINC02605, and CCL5. The microarray datasets used in the study were GSE13896, GSE112260, GSE130928, and GSE16972. In study 1 (GSE13896), a total of 404 dysregulated genes, including 185 upregulated and 219 downregulated genes, were identified in COPD-positive samples. In study 2 (GSE112260), out of a total of 178 DEGs, 50 genes were found to be overexpressed and 128 genes were underexpressed. Study 3 (GSE130928) revealed a total of 350 dysregulated genes, with 168 genes being overexpressed and 182 genes being underexpressed. Additionally, study 4 (GSE16972) identified 36 significantly upregulated and 69 significantly downregulated genes out of a total of 105 dysregulated genes. ADGRE1 and CCL20 were the two common genes identified within these four microarray studies.

The results of the RNA-seq meta-analysis revealed a total of 104 dysregulated genes, with 78 genes being upregulated and 26 genes being downregulated. Among the top 10 upregulated genes were SIGLECI6, ENPP3, BP1FA1, ADAMTS1, STAP1, VGLL3, ITIH5, SCGB1A1, RNU1-28P, and LIN. Conversely, the top 10 downregulated genes included PSPHP1, SOCS3, SLAMF9, CD1A, C20orf96, HLA-H, ADGRG3, SEMA6B, MMP1, and UCHL. Whereas through meta-analysis of microarray studies, a total of 57 differentially expressed genes were identified, consisting of 18 upregulated genes and 39 downregulated genes. The top 10 dysregulated genes included SNAR-D, APAP1L1, FBXO15, SLCM39A10, TMEM163, SRPX, TRERF1, SH3RF1, GPR34 (upregulated genes), and LINCO2244, RSPO3, SHROOM3, ELOVL7, CXCL9, SERPINB, GCH1, TNFAIP6, CCL23, and LAMB1 (downregulated genes).

The GO molecular function analysis of RNA-seq datasets (GSE124180 and GSE183973) in the meta-analysis showed significant upregulation of various molecular functions. These included CXCR3 chemokine receptor binding, CXCR chemokine receptor binding, C-acyltransferase activity, CoA-ligase activity, and fatty-acyl-CoA synthase activity. Of particular interest, CXCR3 and CXCR are known to play crucial roles in immune response and inflammation. The upregulation of these molecules suggests an active immune response, which is often associated with inflammatory processes. The dysregulation of these molecular functions in the context of COPD may reflect the involvement of immune-mediated mechanisms in the disease pathology [37].

The chemokine receptor CXCR3 has been identified as a key determinant of immune responses within secondary lymphoid structures and is known to be highly expressed in Th1 lymphocytes. Its involvement has been linked to the development of COPD. However, recent findings have revealed that chemokine receptors are also expressed on various lung resident cells, including epithelial and smooth muscle cells, raising new questions about their role in the pathogenesis of COPD [38]. The meta-analysis results shed light on the upregulation of fatty-acyl-CoA synthase activity, C-acyltransferase activity, and CoA ligase acidity. These findings suggest an increased utilization of fatty acids as a source of energy and highlight the importance of lipid signaling pathways in the disease. Additionally, the observed upregulation of cholesterol levels may be influenced by smoking, which has a complex impact on lipoprotein metabolism The GO MF analysis of the dysregulated genes in COPD revealed downregulation of several molecular functions, including oxidoreduction-driven active transmembrane, ubiquitin-protein ligase inhibitor activity, and NADH dehydrogenase. These findings suggest potential suppression of these molecular processes in COPD. The downregulation of RNA and rRNA binding reflects a decrease in translational activity, which could impact protein synthesis and overall cellular function. The downregulation of NADH dehydrogenase, an enzyme involved in cellular respiration and oxidoreductase activity, indicates alterations in cellular energy metabolism in the context of COPD. This dysregulation may have implications for the efficiency of energy production and overall cellular respiration. Furthermore, the downregulation of ubiquitin-protein ligase inhibitor activity suggests a potential disruption in the regulation of protein degradation pathways. Ubiquitin-protein ligase inhibitors play a role in controlling the degradation of specific proteins, and their downregulation may contribute to altered protein turnover and homeostasis within the cells [39]. The downregulation of ubiquitin ligase inhibitor activity in COPD indicates potential changes in protein degradation processes. This dysregulation could impact the turnover and stability of proteins within cells, potentially leading to altered cellular functions and protein homeostasis. GO cellular component analysis revealed that the upregulated genes in COPD were being expressed in intracellular vesicles, HFE-transferrin receptor complexes, and membrane attack complexes. The upregulation of genes associated with intracellular vesicles suggests potential changes in cellular processes such as secretion, protein trafficking, and endocytosis. These alterations in intracellular vesicle dynamics may have implications for the overall cellular function and signaling in COPD. Furthermore, the upregulation of HFE-transferrin receptor complexes, which are involved in iron metabolism, suggests potential alterations in iron homeostasis and metabolism in COPD. Iron plays a crucial role in various cellular processes, including oxygen transport, energy production, and oxidative stress. Dysregulation of iron metabolism could contribute to the pathogenesis of COPD and its associated complications [40].

The GO MF meta-analysis of microarray datasets revealed the upregulation of syndecan binding, indicating an important role for syndecans in cell adhesion and signaling. Syndecans are cell surface proteoglycans that play a critical role in mediating cell-cell and cell-matrix interactions, as well as modulating various signaling pathways. The observed upregulation of syndecan binding suggests that these molecules may be involved in enhanced cell adhesion and communication processes in the context of the studied condition [41]. In individuals with COPD, syndecan serves as an inflammatory biomarker associated with lung function and systemic inflammation. Its involvement in the inflammatory process is significant as it interacts with ligands such as chemokines and cytokines. The upregulation of syndecan bindings facilitates cellular interactions and modulates signaling pathways. Furthermore, the upregulated endodeoxyribonuclease activity plays a role in DNA metabolism, specifically in repairing damage caused by reactive oxygen species, which includes oxidized bases, single-strand breaks, and abasic sites. However, it is worth noting that the increased DNA damage observed in COPD lungs, including double-strand breaks, is unlikely to be solely attributable to cigarette smoke. Notably, the lungs of COPD patients exhibit higher levels of DNA damage, indicating factors beyond smoking contribute to the observed DNA damage, distinguishing COPD cases from smokers and non-smokers without the condition [42]. The G protein-coupled purinergic nucleotide was identified as being upregulated and playing a role in signal transmission across the cell membrane. This process involves the activation of associated G proteins to initiate downstream signaling events [43]. Nevertheless, in individuals with COPD, G protein-coupled receptors participate in the development of airflow limitation and airway hyperresponsiveness, both of which are key factors in the pathophysiology of COPD. The relaxation of airway smooth muscle, achieved through the activation of β2-adrenergic receptors and the inhibition of muscarinic receptors, is crucial in the management of COPD symptoms [44]. Additionally, elevated levels of MAP-kinase scaffold activity and purinergic nucleotide receptor activity were detected. MAP-kinases serve as scaffolds that enable cells to interpret external signals, particularly those associated with inflammation and cellular responses to stress [45].

The GO CC analysis provided intriguing insights into COPD. The upregulated genes were found to be expressed in various cellular components, including the piP-body, integral component of synaptic vesicles, P granules, AMPA glutamate receptor complexes, and intrinsic components of synaptic vesicles. These components are primarily associated with synaptic vesicles and synaptic activity in neurons. These findings suggest the possibility of alterations in synaptic function and signaling in the context of COPD. Synaptic vesicles play a crucial role in several pulmonary functions. Notably, serotonin, a neurotransmitter involved in pulmonary functions, has been associated with oxidative stress, which is known to contribute to the pathogenesis of COPD. Furthermore, dysregulation of synaptic vesicle function can lead to functional impairments, and it is plausible that COPD may be associated with impaired synaptic vesicle function, which could potentially contribute to the development of the disease [46].

Additionally, the downregulated genes exhibited enrichment in various cellular components, including the lumenal side of the endoplasmic reticulum, luminal side of the endoplasmic reticulum, luminal side of the endoplasmic reticulum, ER to Golgi transport vesicle membrane, coated vesicle membrane, and mitochondrial envelope. Conversely, the downregulated genes associated with COPD were found to be enriched in cellular components related to the endoplasmic reticulum, integral component of the luminal side of the endoplasmic reticulum, ER to Golgi transport vesicle membrane, and coated vesicle membrane. The downregulation of these cellular components indicates potential disruptions in endoplasmic reticulum homeostasis and protein trafficking processes in the context of COPD [47]. The enrichment analysis of gene ontology biological processes revealed that the upregulated genes were significantly associated with positive regulation of mitochondrial depolarization and negative regulation of lens fiber cell differentiation. Furthermore, the downregulated genes exhibited enrichment in various biological processes, including cytokine-mediated signaling, response to lipopolysaccharide, and regulation of cytokine production. These findings suggest an altered immune response in the context of COPD.

Furthermore, the meta-analysis of RNA-seq data using the KEGG pathway database identified several upregulated pathways, including the Toll-like receptor signaling pathway, butanoate metabolism, cytokine-cytokine receptor interaction, and chemokine signaling pathway. Toll-like receptors (TLRs) are membrane-bound receptors that play crucial roles in the innate immune system and have been implicated in the pathogenesis of COPD. The chemokine signaling pathway, on the other hand, is involved in regulating leukocyte migration during development and inflammation. Dysregulation of chemokines, which are signaling molecules involved in directing cell migration, has been associated with various inflammatory diseases. Interestingly, the KEGG pathway meta-analysis of microarray revealed that lysosome was highly upregulated in COPD. The key function of lysosomes is digestion and removal of waste while dysfunctional lysosomes alter protein degradation in COPD [48]. In contrast, pathways associated with immune cell differentiation, including Th1 and Th2 cellular differentiation, Th17 cell differentiation, and C-type signaling pathway, were observed to be downregulated in COPD. These pathways play significant roles in immune responses and the regulation of inflammation. The downregulation of these pathways suggests a potential impairment in immune responses and inflammatory regulation in COPD.

The meta-analysis of RNA-seq and microarray datasets in relation to COPD revealed the presence of seven common genes. These genes, namely VGLL3, ITIH5, ELOVL7, ACOD1, LAMB1, CXCL9, and GBP5, were consistently identified across both types of analyses. The identification of these common genes highlights their potential relevance in the context of COPD and suggests their involvement in the molecular mechanisms underlying the disease. Vestigial-like family member 3 (VGLL3) is involved in several cellular processes, such as cell proliferation and differentiation [49]. However, VGLL3 has been involved in respiratory conditions such as COPD [50]. On the other hand, Inter-Alpha-Trypsin Inhibitor Heavy Chain 5 (ITIH5) is involved in inflammatory processes and immune system regulation which are known to be involved in COPD development [51]. Elongation of very long chain fatty acids protein 7 (ELOVL7) is involved in the synthesis of fatty acids [52] which have a pathological connection to COPD. Moreover, It has been demonstrated that Aconitate Decarboxylase 1 (ACOD1) is involved in the control of oxidative stress and inflammation [53] which are key mechanisms in COPD pathogenesis. The extracellular matrix substance Laminin Subunit Beta 1 LAMB1 has been associated with lung tissue remodeling and healing [54]. CXCL9 is involved in immune cell recruitment [55] while the immune response to viral infections is aided by Guanylate Binding Protein 5 (GBP5) [56].

In this study, a comprehensive approach was employed to identify key genes and protein-protein interactions associated with the pathology of COPD. Dysregulated genes obtained from both microarray and RNA-seq meta-analysis experiments were subjected to protein-protein interaction analysis. Based on the degree of centrality and the number of interactions, the top 10 hub genes were identified. Among these, a common gene that exhibited the highest number of interactions with other proteins in the network, including both microarray and RNA-seq studies, was selected for further analysis in the context of drug repurposing. The focus of the investigation was on the CXCL9 gene, which emerged as a prominent hub gene with high interactivity. CXCL9 has been implicated in the pathogenesis of COPD and plays a crucial role in the disease. To assess the feasibility of targeting CXCL9, the 3D structure of its protein was subjected to molecular docking analysis with a panel of known drug compounds. These compounds have either been experimentally validated or are currently undergoing clinical trials for the treatment of various lung diseases and cancer. The main objective of this study was to evaluate the potential of inhibiting CXCL9, which is upregulated in COPD patients, using these drug compounds. This analysis aimed to provide insights into the possibility of repurposing existing drugs to target CXCL9 and potentially mitigate the pathology of COPD. Cancer is a leading cause of disease worldwide that involves uncontrolled division of abnormal cells due to several modifications in gene expression [57]. In 2020, nearly 10 million deaths were accounted globally that comprised a major portion of breast cancer, lung cancer, colon and rectum cancer, prostate cancer, skin cancer and stomach cancer [58]. Out of all the types of cancers, lung cancer is the leading cause of deaths in which the major cause is chronic smoking in addition to other contributing factors [59]. A compound, nintedanib, has been approved for the treatment of Idiopathic Pulmonary Fibrosis (IPF), a progressive and fatal lung cancer [60]. Another compound, tepotinib is a Tyrosine Kinase Inhibitor (TKI) developed mainly for the treatment of NSCLC patients [61].

Moreover, the molecular docking results indicate that CXCL9-Nintedanib was the best-docked complex among all the others, followed by CXCL9-Tepotinib and CXCL9-Crizotinib based on their docking scores, and was observed to have the binding scores of −7.3 (kcal/mol), −6.8 (kcal/mol) and −6.2 (kcal/mol) respectively. Overall, the best complex was CXCL9-Nintedanib, in which nintedanib interacted with two residues (LYS-103 and GLN-106) of CXCL9 protein (Fig 4**(b)****(c)**). Furthermore, nintedanib, an intracellular tyrosine kinase inhibitor (TKI) has recently been approved for use in chronic fibrosing interstitial lung diseases (ILDs) and systemic sclerosis-associated ILD (SSc-ILD). Moreover, it was also one of the first drugs approved for use in idiopathic pulmonary fibrosis (IPF) [62]. Finally, the molecular dynamics simulation results indicated that the protein, CXCL9, showed considerable instability when docked with all the selected compounds (Nintedanib, Tepotinib, and Crizotinib). However, the CXCL9 protein overall showed the highest fluctuations (RMSD: 22.79 Å and RMSF: 20.27 Å) with the Crizotinib (Fig 5). Moreover, the CXCL9 protein showed the best protein-ligand and ligand-protein contacts with Nintedanib as it showed the most considerable interactions with the CXCL9 protein (Fig 6). On the other hand, all the ligands (Nintedanib, Tepotinib, and Crizotinib) fluctuated considerably, as well, when docked with CXCL9 protein. Although, crizotinib showed good simulation results with the CXCL9 protein but its binding affinity [-6.2 (kcal/mol)] was not good enough as of the CXCL9-Nintedanib complex [-7.3 (kcal/mol)]. So, ultimately the best docked complex for the inhibition of CXCL9 protein was CXCL9-Nintedanib complex.

In addition, if compounds such as nintedanib, tepotinib, and crizotinib have demonstrated potential in combating various types of cancers, there is a possibility that they could be repurposed for targeting the CXCL9 protein. Nintedanib, in particular, could be investigated for its efficacy against CXCL9 due to its known properties and effects on different cancers. Utilizing already approved drugs for a new purpose offers the advantage of having existing knowledge about their safety, pharmacokinetics, and toxicity profiles. Repurposing existing drugs can also save time and resources compared to developing entirely new drugs.

However, further research is necessary to fully comprehend the interactions between nintedanib and the CXCL9 protein for therapeutic purposes. This would involve studying the binding affinity, functional consequences, and potential downstream effects of the interaction. By gaining a deeper understanding of these interactions, researchers can evaluate the feasibility of repurposing nintedanib or other related compounds for targeting CXCL9 as a potential therapeutic strategy for COPD.

## Conclusion

Drug reprofiling for diseases that show resistance to the current therapeutics has offered the benefits of liberating time and resources by utilizing the existing knowledge of their toxicity, pharmacokinetics, and safety. In this study, the hub gene, CXCL9 which is a chemokine, significantly associated with inflammation and immune response was shortlisted as a novel therapeutic target after a comprehensive meta-analysis of differentially expressed genes identified via multiple microarray and RNA-seq datasets of healthy control vs. diseased patients from alveolar macrophage in COPD. Significantly enriched pathways and GO terms elaborated on the effectiveness of CXCL9 to be utilized as a novel target for the treatment of COPD. Moreover, virtual screening of CXCL9 against the selected FDA-approved drugs revealed that Nintedanib, Tepotinib, and Crizotinib were the top three drugs showing better binding affinity to inhibit CXCL9. Furthermore, molecular dynamics simulations disclosed better results of CXCL9-Crizotinib as compared to Nintedanib. However, based on the docking results effective inhibition of CXCL9 might be exhibited by Nintedanib as it showed the highest scores. Further functional downstream analysis and experimental validations are mandatory to establish Ninatedanib as a therapeutic agent for COPD treatment.

## Supporting information

Supplementary Sheet 1

## Acknowledgements

The author pays gratitude to the Teesside University, school of health and life sciences (SHLS).

