## Supplementary Document 1 for "Exploring Inflammatory Dysregulation in Alveolar Macrophages: Implications for Novel Therapeutic Targets in Chronic Obstructive Pulmonary Disease"

### Gene ontologies and KEGG pathways enrichment for RNA-seq and microarray-based DEGs through EnrichR

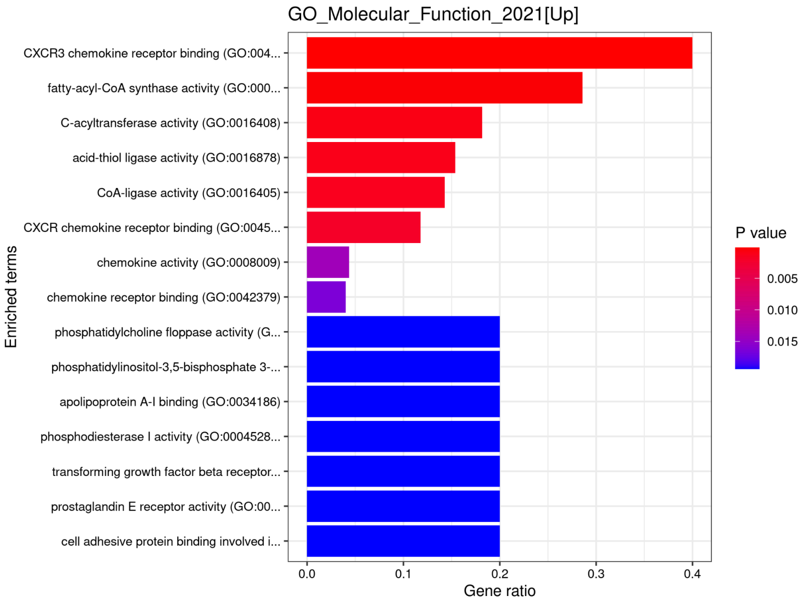


**Figure S1.** Upregulated molecular functions gene ontologies identified through EnrichR.

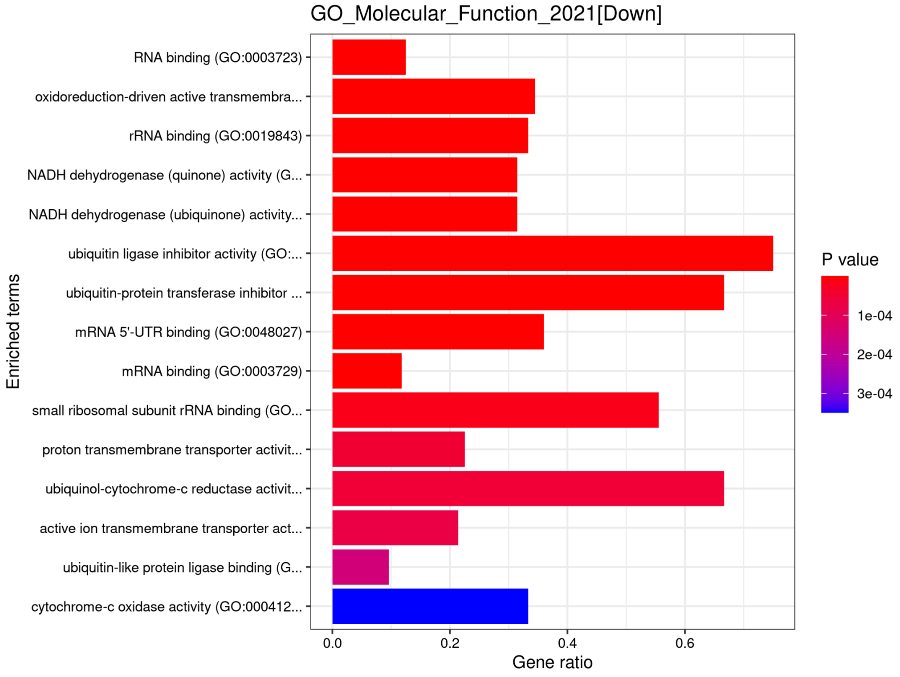


**Figure S2.** Downregulated molecular functions gene ontologies identified through EnrichR.

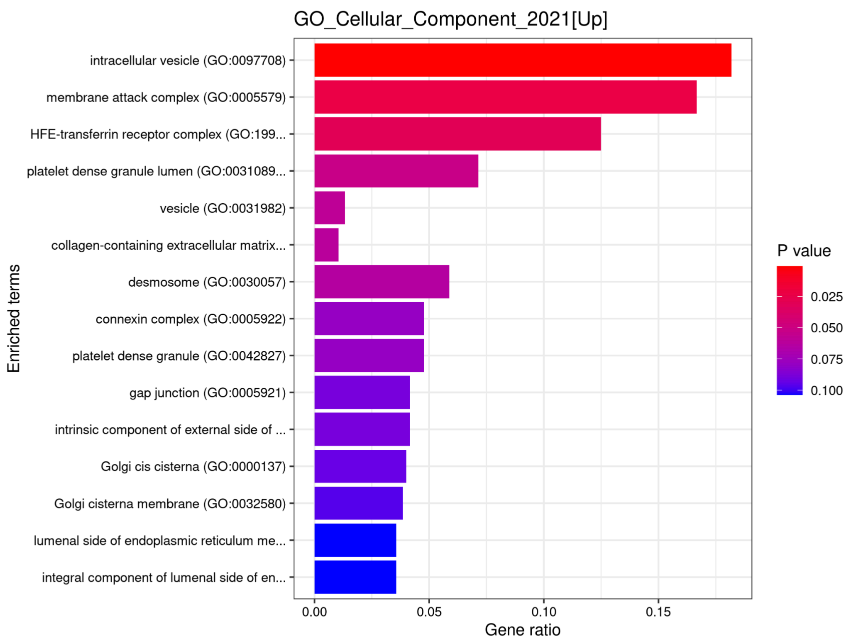


**Figure S3.** Upregulated cellular component gene ontologies identified through EnrichR.

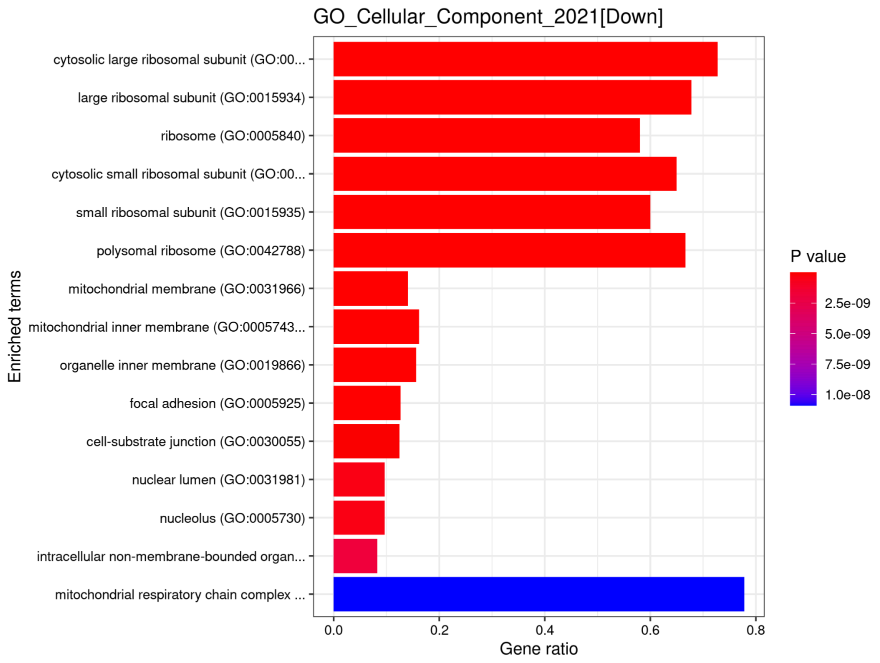


**Figure S4.** Downregulated cellular component gene ontologies identified through EnrichR.

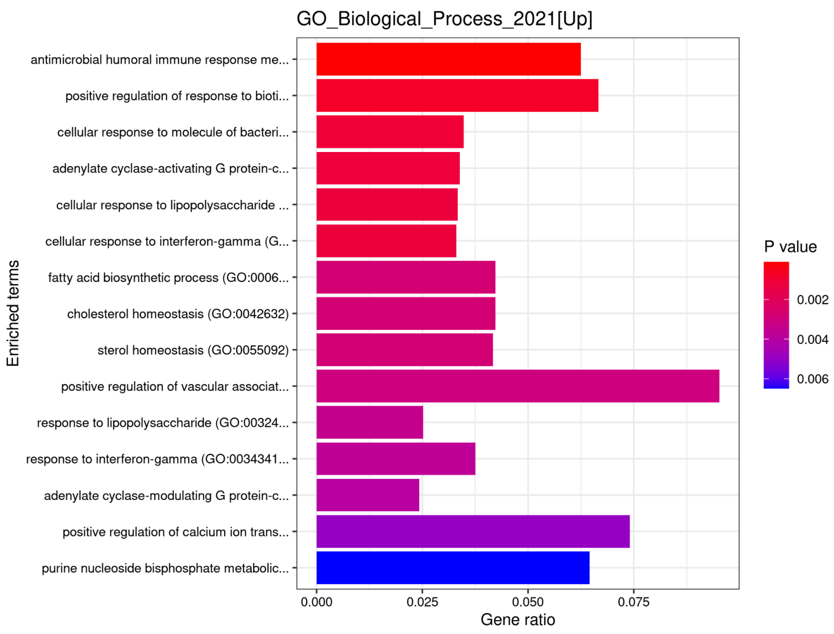


**Figure S5.** Upregulated biological processes gene ontologies identified through EnrichR.

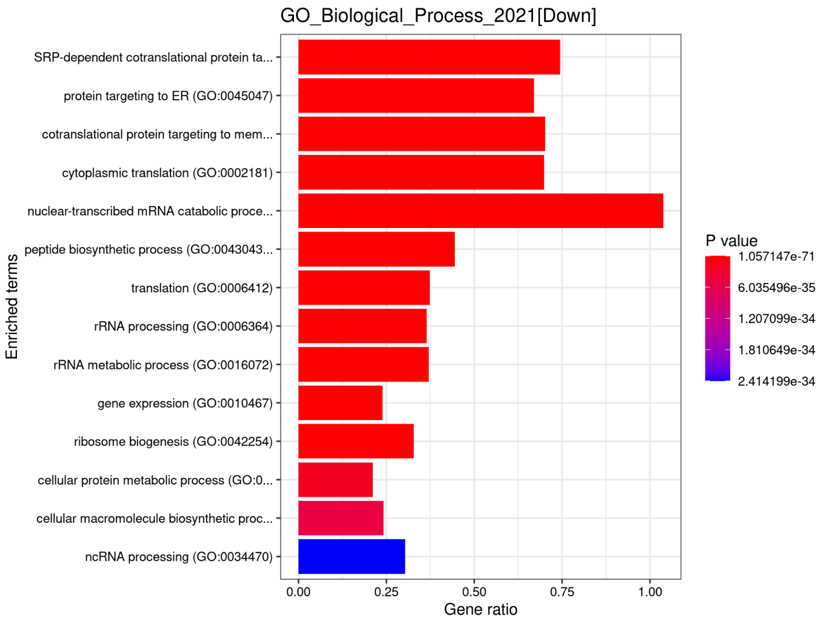


**Figure S6.** Downregulated biological processes gene ontologies identified through EnrichR.

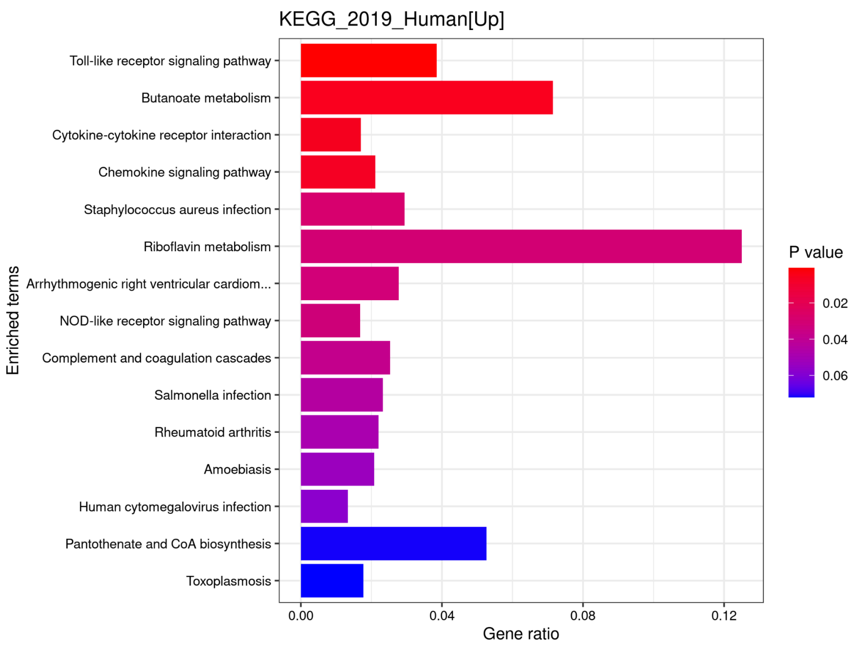


**Figure S7.** Upregulated KEGG biological pathways identified through EnrichR.

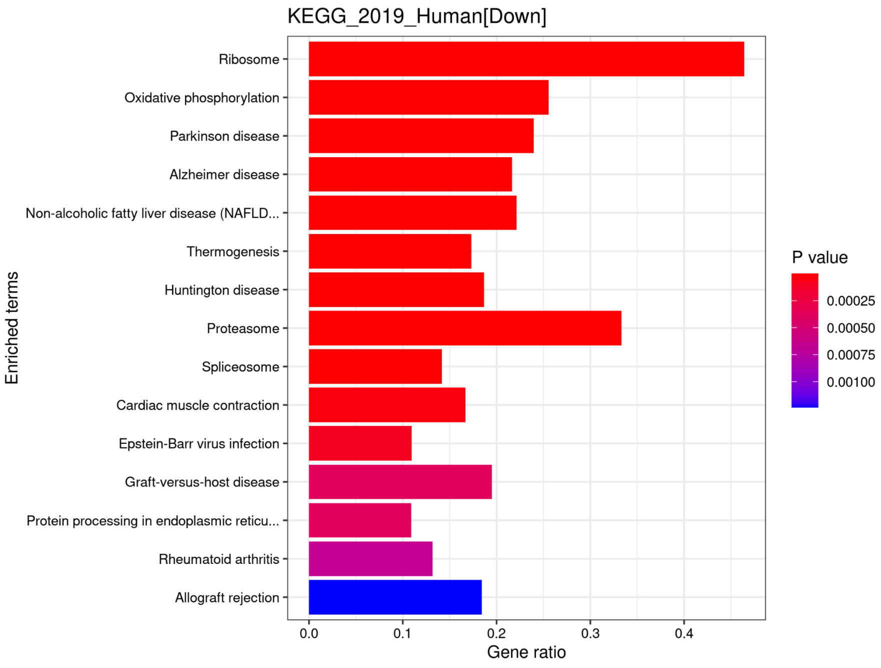


**Figure S8.** Downregulated KEGG biological pathways identified through EnrichR.

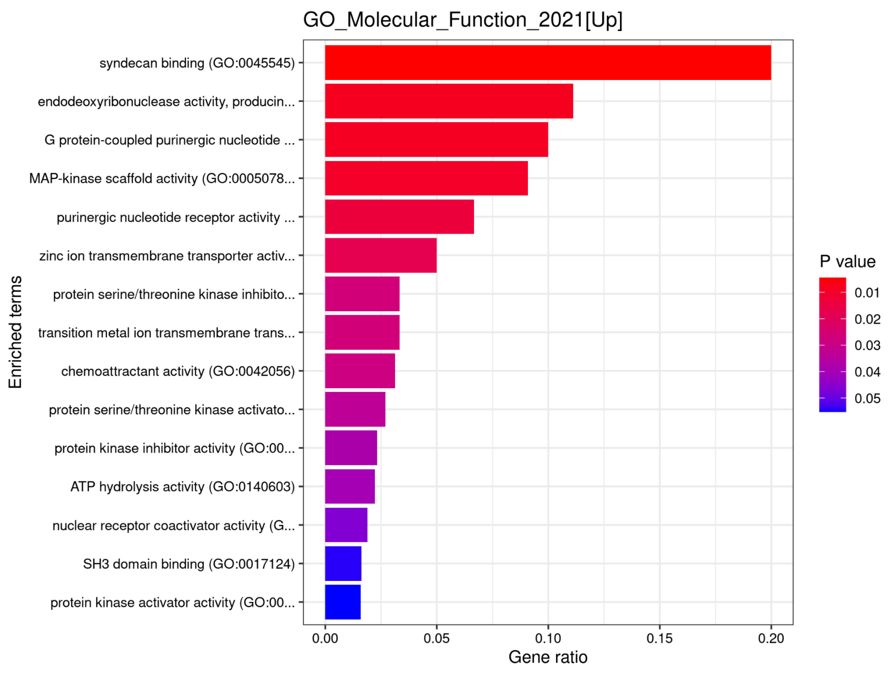


**Figure S9.** Upregulated molecular functions gene ontologies identified through EnrichR.

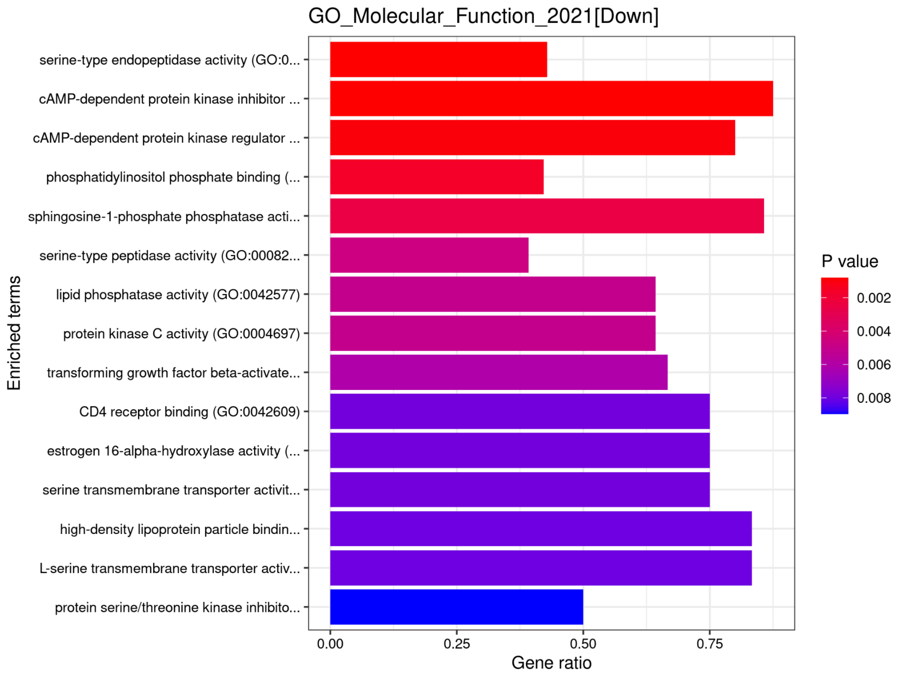


**Figure S10.** Downregulated molecular functions gene ontologies identified through EnrichR.

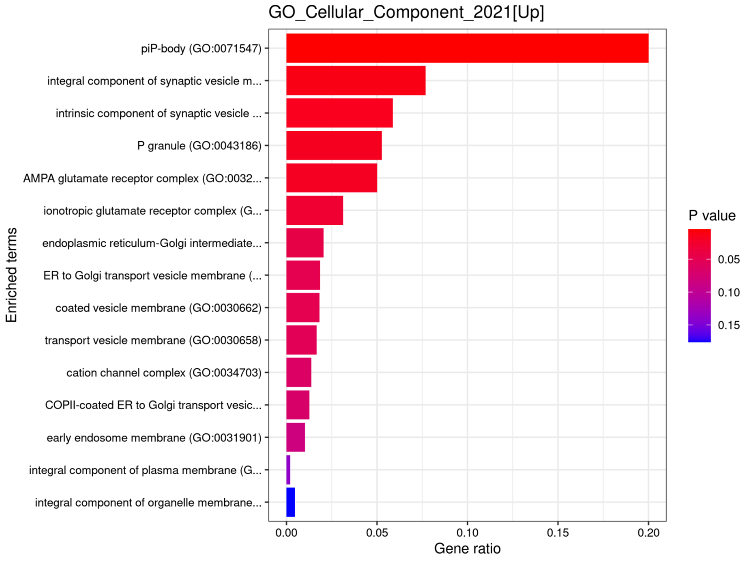


**Figure S11.** Upregulated cellular components gene ontologies identified through EnrichR.

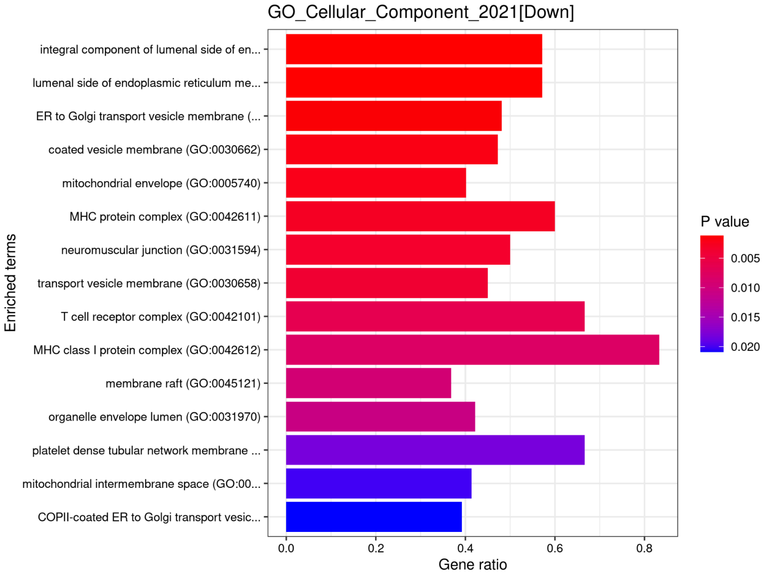


**Figure S12.** Downregulated cellular components gene ontologies identified through EnrichR.

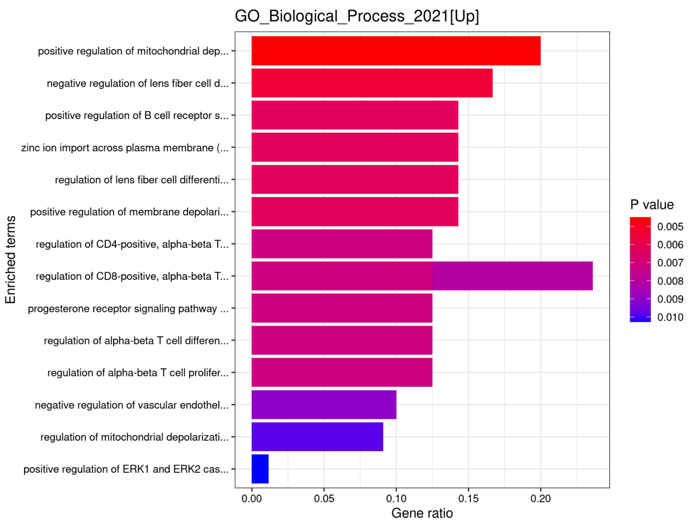


**Figure S13.** Upregulated biological processes gene ontologies identified through EnrichR.

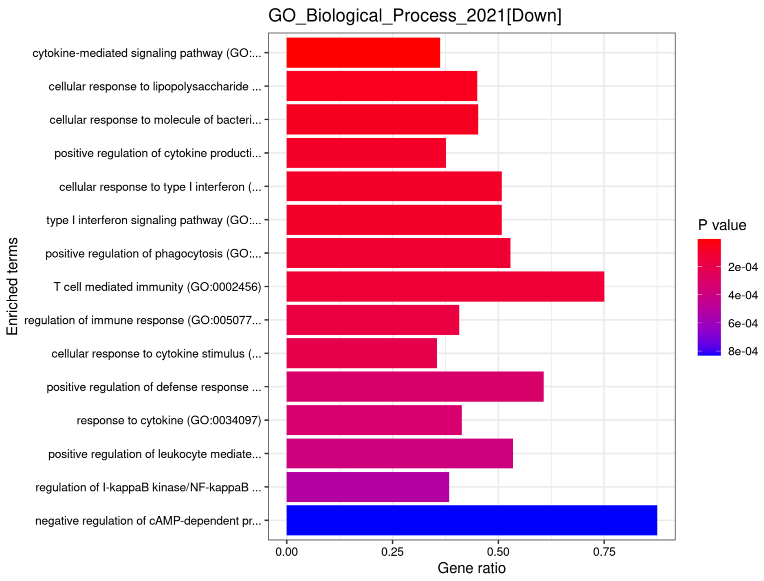


**Figure S14.** Downregulated biological processes gene ontologies identified through EnrichR.

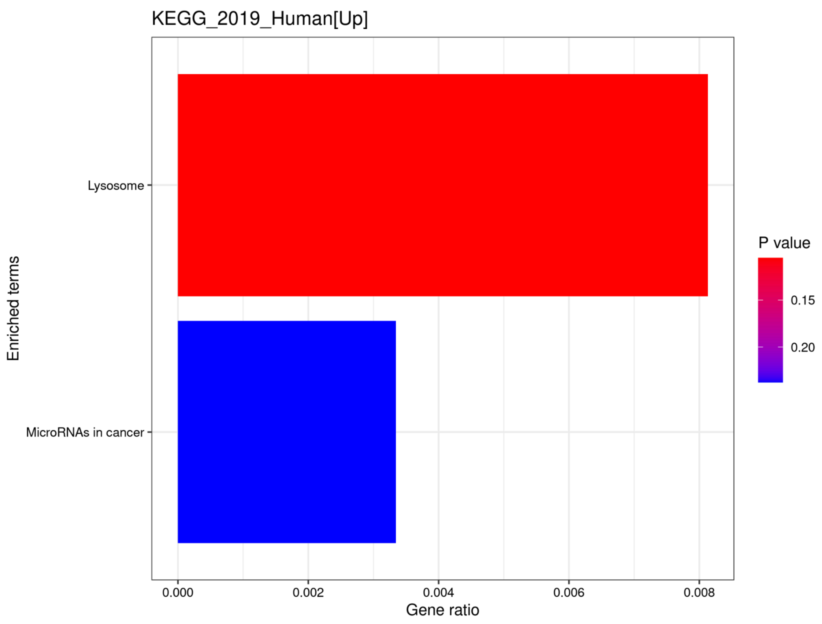


**Figure S15.** Upregulated KEGG biological pathways identified through EnrichR.

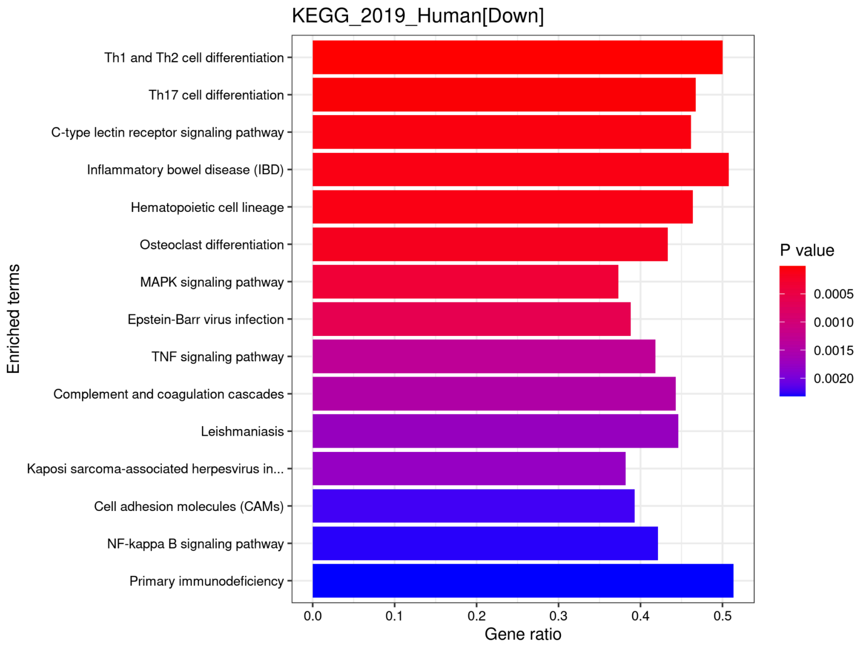


**Figure S16.** Downregulated KEGG biological pathways identified through EnrichR.

### Protein-Protein Interaction Analysis of RNA-seq and microarray-based DEGs through STRING

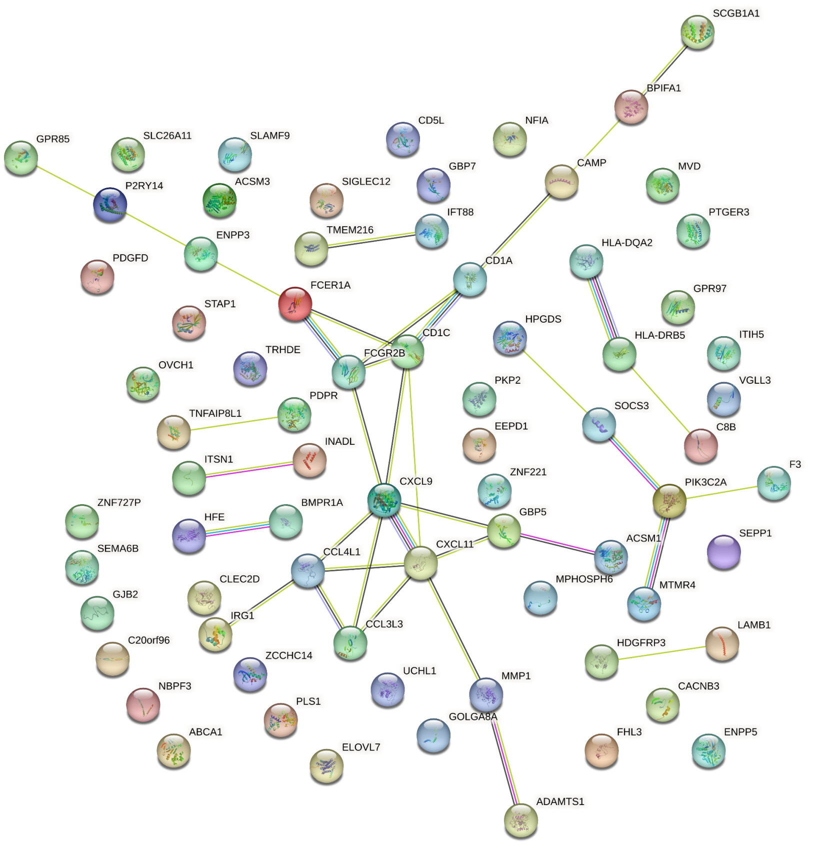


**Figure S17.** PPIs of the dysregulated meta-analysis genes based on RNA-seq studies.

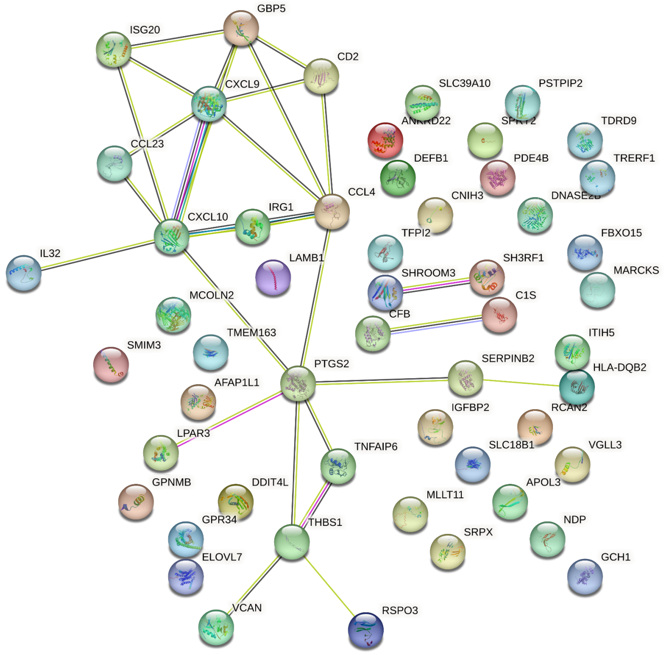


**Figure S18.** PPIs of the dysregulated meta-analysis genes based on microarray studies.
